# Perinatal Brain Injury Triggers Niche-Specific Changes to Cellular Biogeography

**DOI:** 10.1101/2024.02.25.581974

**Authors:** Nareh Tahmasian, Min Yi Feng, Keon Arbabi, Bianca Rusu, Wuxinhao Cao, Bharti Kukreja, Asael Lubotzky, Michael Wainberg, Shreejoy J. Tripathy, Brian T. Kalish

## Abstract

Preterm infants are at risk for brain injury and neurodevelopmental impairment due, in part, to white matter injury following chronic hypoxia exposure. However, the precise molecular mechanisms by which perinatal hypoxia disrupts early neurodevelopment are poorly understood. Here, we constructed a brain-wide map of the regenerative response to newborn brain injury using high-resolution imaging-based spatial transcriptomics to analyze over 1.3 million cells in a mouse model of chronic neonatal hypoxia. Additionally, we developed a new method for inferring condition-associated differences in cell type spatial proximity, enabling the identification of niche-specific changes in cellular architecture. We observed hypoxia-associated changes in region-specific cell states, cell type composition, and spatial organization. Importantly, our analysis revealed mechanisms underlying reparative neurogenesis and gliogenesis, while also nominating pathways that may impede circuit rewiring following perinatal hypoxia. Altogether, our work provides a comprehensive description of the molecular response to newborn brain injury.

## INTRODUCTION

Preterm birth is the leading cause of perinatal morbidity and mortality in developed countries and affects approximately 15 million infants annually.^1^ While significant strides have been made in perinatal health care leading to increased survival rates, the long-term impact of premature birth on the trajectory of neurodevelopment remains a major unsolved challenge. Survivors of preterm birth are at high risk for neurodevelopmental impairment, including cognitive and motor disability, as well as autism spectrum disorder, epilepsy, and attention-deficit/hyperactivity disorder.^2–5^ The causes of brain injury in preterm infants are complex and multifactorial, but chronic exposure to hypoxia (HX) due to lung immaturity is an important driver of brain injury and maldevelopment. Importantly, white matter injury (WMI) is most prevalent in infants born at early gestational ages preceding the onset of myelination. This suggests a developmental susceptibility of immature oligodendrocyte (OL)-lineage cells to HX-ischemia, oxidative stress, and impaired cerebral perfusion, leading to maturational failure.^6^ Despite this understanding, we lack therapies to protect the preterm brain or to promote recovery during this critical developmental window.

A mouse model of chronic sublethal HX, in which neonatal mice are exposed to one week of 10% oxygen, replicates many structural and pathophysiological aspects of human preterm injury and has been widely used to study preterm WMI.^6–9^ However, most studies have focused on injury and repair within the OL-lineage, rather than brain-wide changes in the response to perinatal HX and the role of cell-cell communication networks. Importantly, there is growing recognition that the local microenvironment in specific brain regions dictates many aspects of cell fate and state^10–12^, which highlights the necessity of characterizing brain region-specific cellular programs that govern recovery and repair. Glial identity varies based on cortical location^13–15^ and the local spatial niche may inform injury response.^16,17^ It is therefore likely, but incompletely understood, that a similar region-specific injury response occurs in the context of chronic neonatal HX.

To address this gap in knowledge, we applied spatially resolved single-cell transcriptomics (multiplexed error-robust fluorescence *in situ* hybridization, MERFISH) to explore how neonatal HX affects the brain’s spatial organization and signaling landscape. Unlike traditional droplet-based single-cell genomics, MERFISH captures sub-cellular localization of gene expression within the native tissue context, offering insight into cellular organization and architecture.^18–20^ We performed MERFISH on the mouse brain following chronic sublethal neonatal HX and on age-matched normal oxygen (normoxic, NX) controls. At baseline (under NX conditions), we identified brain region-specific cell states and signaling-associated gene expression profiles in non-neuronal cells. Following neonatal HX, we identified increased oligodendrocyte precursor cell (OPC) proliferation but arrested differentiation of these cells into mature OL in the subventricular zone (SVZ) and corpus callosum. We characterized region- and cell type-specific gene expression changes that may inhibit functional myelination and repair, including upregulation of extracellular matrix components and other mediators known to inhibit oligodendrogenesis. Further, to quantify compositional changes to the cellular microenvironment after HX, we developed a novel approach to infer changes in the spatial proximity of neighboring cell types. Using this method, we found extensive reorganization of local cellular architecture in the cortex, SVZ, and corpus callosum following HX. Overall, these results provide novel insights into the mechanisms underlying white matter regeneration in the neonatal brain and underscore the complex multicellular processes in the regenerative response to HX-induced preterm WMI.

## RESULTS

We employed a widely used mouse model of chronic sublethal HX to investigate the brain’s regenerative response to perinatal injury (Figure 1A, experimental schema). Mouse litters were exposed to 10% oxygen (HX) or 21% oxygen (normoxia, NX) between P3-11 (see *Methods*), a developmental period in mice that resembles the third gestational trimester in humans.^21^ Mouse dams were rotated daily between normoxia and hypoxia conditions to avoid any confounding based on maternal care or nutrition. All mice were then moved to an environment with normal levels of oxygen until they were sacrificed at P21 (see *Methods*). We focused specifically on P21 as it represents a peak time during the post-injury repair period.

**Figure 1:**
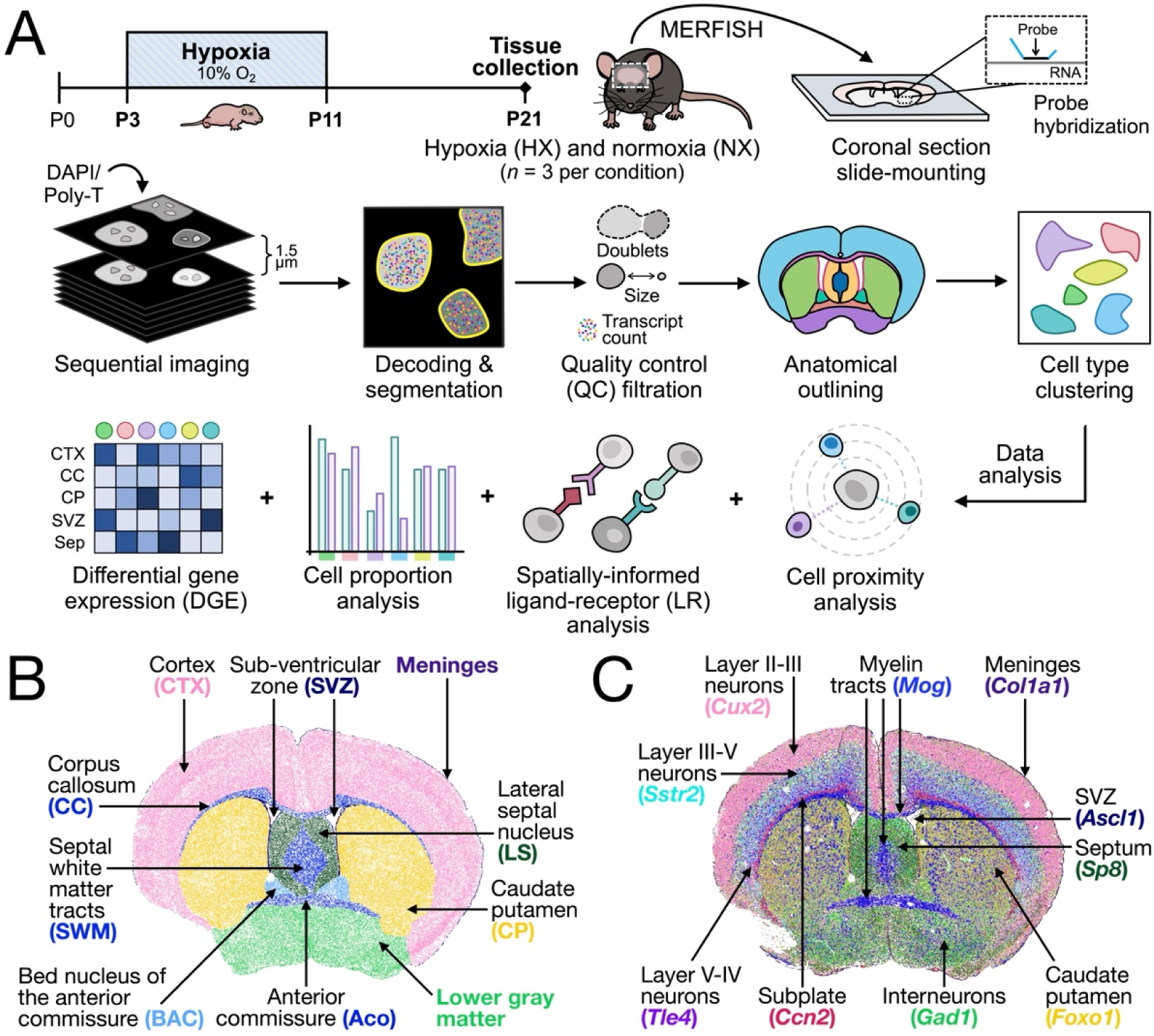
Multiplexed error-robust fluorescent *in situ* hybridization (MERFISH) sequencing identifies all major cell types and anatomical regions of the postnatal murine brain. **(A)** Schematic representation of the HX paradigm, tissue harvesting, tissue processing, quality control, and downstream data analysis approach applied to the MERFISH dataset. Tissue was harvested and processed for postnatal day 21 (P21) HX and NX mice (*n* = 3 male mice per condition). MERFISH-derived spatial map displaying the **(B)** major anatomical regions outlined during sample processing steps, and **(C)** a select set of genes used to identify and outline the anatomical regions utilized in downstream data analysis. Areas of the brain are colored according to their anatomical region or major identifying gene.

We performed multiplexed error-robust fluorescence *in situ* hybridization (MERFISH) on coronal brain sections from three HX and three NX mice (see *Methods*). An RNA probe panel of 497 genes (Table S1) was selected to enable the identification of major forebrain cell types as well as components of biologically relevant pathways including growth factors, Wingless-related integration site (WNT) /LJ-catenin, bone morphogenic protein (BMP), immune recognition and regulation, perineuronal net components, and senescence-associated genes (see *Methods*). Cells were segmented using a deep-learning algorithm, CellPose 2.0^22^, to segment cells based on DAPI, a nuclear stain, and PolyT, an mRNA stain (see *Methods*). After segmentation, transcript counts were normalized by volume and quality control filtering was performed to remove segmented cells with abnormal physical size, low total number of transcripts, and potential doublets (Figure 1A, see *Methods*). The final dataset, encompassing all replicates, profiled 1,380,953 cells which had a median diameter of 179.5 µm and a median expression of 285.6 genes per cell.

Each coronal MERFISH image of the mouse brain was labeled based on region-specific marker genes and anatomic landmarks to define regions for downstream analysis (Figures 1B-C). After normalizing each sample, we independently integrated and analyzed cells in the cortex, corpus callosum, SVZ, caudate putamen (CP), lateral septal nucleus (LS), anterior commissure (ACo), and septal white matter tracts (SWM), respectively, across all datasets. Highly variable genes were used to compute principal components for uniform manifold approximation and projection (UMAP) and shared nearest neighbor clustering. After filtering, the final dataset included 668,947 cortex, 8,708 corpus callosum, 10,754 SVZ, 83,235 CP, 19,088 LS, 4,311 ACo, and 12,837 SWM cells across all samples (Figures 2A-B, 3A, 3E, and S2A-D). Our analyses focused on the cortex, corpus callosum, and SVZ, given their relevance to HX injury and ample cell numbers, contributing statistical power to our findings. The number of unique genes and transcripts per cell for these three regions are displayed in Figures S1A-F.

**Figure 2:**
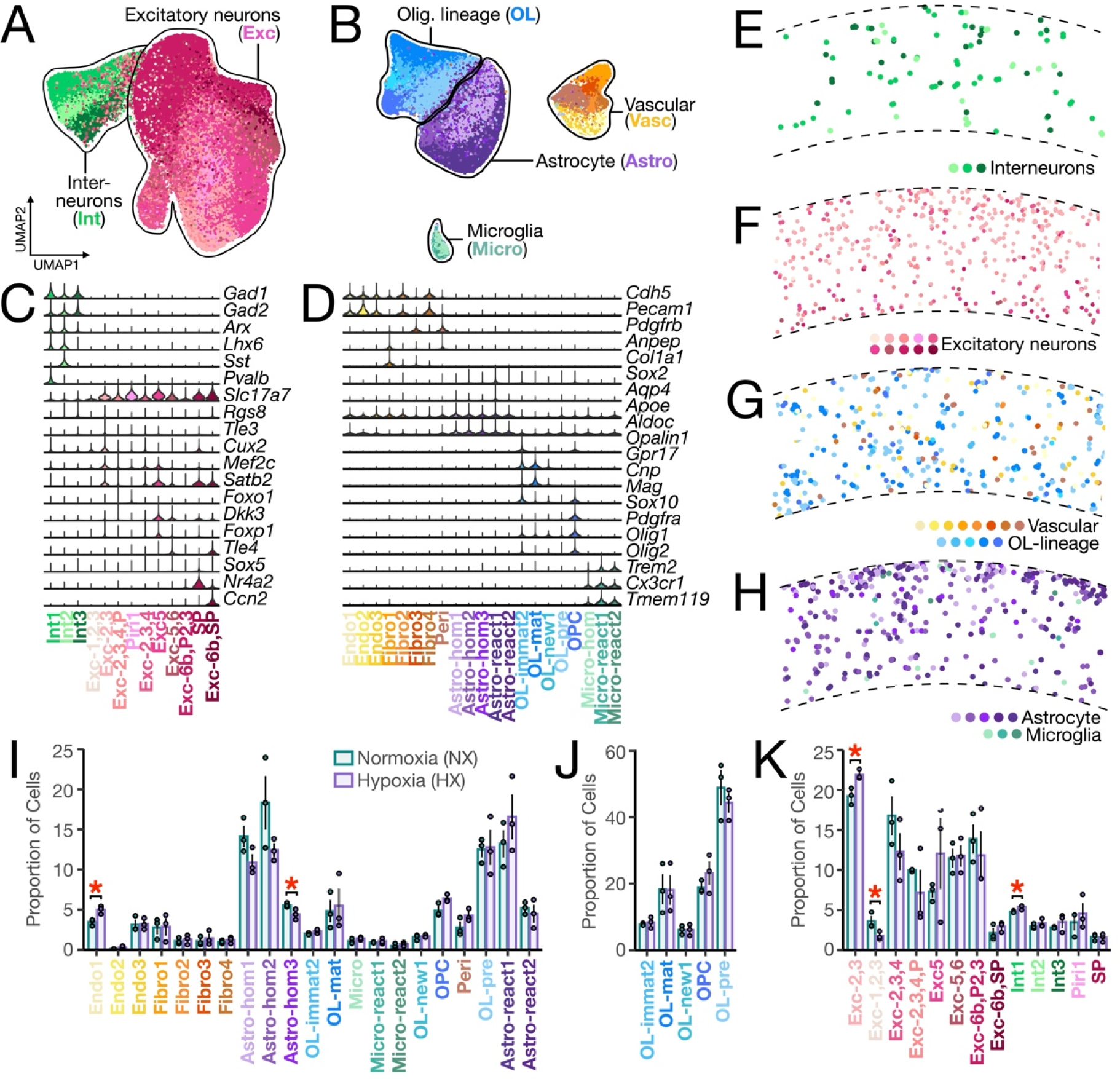
Cell type classification and spatial distribution in the P21 mouse cortex. UMAP visualization of cortical **(A)** neuronal and **(B)** non-neuronal cells where each dot represents a single cell, for a total of 668,947 cells. UMAP plots are generated from combined replicates across NX and HX conditions. Each cluster is colored by Cell Subtype and clusters are outlined and labelled by Cell Type. Violin plots showing gene expression patterns of **(C)** neuronal and **(D)** non-neuronal Cell Subtypes identified in the cortex with several key canonical marker genes used for cluster identification. The width of the violin corresponds to the proportion of nuclei expressing the indicated gene, and the color of the violin corresponds to the Cell Subtype as follows: Exc: excitatory neuron, Int: interneuron, Exc-1,2,3: cortical layers I-III excitatory neurons, Exc-2,3: cortical layers II-III excitatory neurons, Exc-2,3,4: cortical layers II-IV excitatory neurons, Exc-2,3,4,P: cortical layers II-IV excitatory neurons and piriform neurons, Exc-5: cortical layer V excitatory neurons, Exc-5,6: cortical layers V-VI excitatory neurons, Exc-6b,P2,3: cortical layer VIb excitatory neurons and piriform neurons, Exc-6b,SP: cortical layer VI excitatory neurons and subplate neurons, Piri: piriform neurons, SP: subplate neurons, Vasc: vascular cell, OL: oligodendrocyte-lineage cell, Astro: astrocyte, Micro: microglia, OL-immat: immature oligodendrocyte, OL-mat: mature oligodendrocyte, OL-new: newly-formed oligodendrocyte, OPC: oligodendrocyte precursor cell, OL-pre: OPC precursor, Astro-hom: homeostatic astrocyte, Astro-react: reactive astrocyte, Micro-hom: homeostatic microglia, Micro-react: reactive microglia, Endo: endothelial cell, Fibro: fibroblast, Peri: pericyte. Spatial plots displaying the distribution of **(E)** interneurons **(F)** excitatory neurons, **(G)** vascular and OL-lineage cells, and **(H)** astrocytes and microglia in the cortex. Bar plots displaying the proportion of **(I)** non-neuronal Cell Subtypes, **(J)** OL-lineage Cell Subtypes, and **(K)** neuronal Cell Subtypes in the cortex of HX versus NX mice. The x-axis displays all major Cell Types and Cell Subtypes profiled through MERFISH. Data obtained from *n*L=L3 male biological replicates per condition. Bars represent the average values for each condition, and dots represent the average values for each mouse per condition. Significance is determined using the two-tailed Student’s t-test. *p-value < 0.05; **p-value < 0.01. Comparisons not labeled with asterisks are not significant. Error bars represent the average ± 1 standard deviation. Each bar is labeled by Cell Subtype as previously defined.

### Cell type identification by MERFISH

A major benefit of high-resolution spatial transcriptomics is the potential to characterize cell type diversity within defined anatomical borders. As such, we analyzed each major anatomic region independently to ascertain region-specific cell type composition and spatially restricted cell types. In the cortex and corpus callosum, we characterized broad cell type categories (from here on referred to as “Cell Types”) including neuronal cells present only in the cortex, corresponding to excitatory neurons and interneurons (Figure 2A); and non-neuronal cells in both regions corresponding to OL-lineage cells, vascular and perivascular cells (from here on collectively termed vascular cells), astrocytes, and microglia (Figure 2B). Excitatory neuron clusters were identified by their expression of *Slc17a7* and *Slc17a6*, interneurons by *Gad1* and *Gad2*, OL-lineage cells by *Olig2*, *Sox8* and *Sox10*, astrocytes by *Aqp4* and *S100b*, vascular cells by *Pecam1* and *Cdh5*, and microglia by *Cx3cr1* and *Tmem119* (Figures 2C-D). Next, each Cell Type group was sub-clustered to identify subtypes or region-specific cell states (the “Cell Subtypes”). From the cortex, excitatory neurons formed 10 clusters which were annotated based on expression of cortical layer-specific markers (Figure S3B). These included a cluster of cells that were spatially concentrated in cortical layers I-III and expressed *Cux2* and *Mef2c* (Exc-1,2,3); a cluster of cells spatially concentrated in cortical layers II and III which expressed *Cux2*, *Mef2c*, and *Satb2* (Exc-2,3); a cluster of cells spatially concentrated in cortical layers II-IV and expressed *Mef2c* and *Lmo4* (Exc-2,3,4); a cluster of cells spatially concentrated in cortical layers II-IV and all three layers of the piriform cortex and expressed *Mef2c*, *Pou3f3,* and *Lmo4* (Exc-2,3,4,P); a cluster of cells spatially concentrated in the piriform layer 1 and expressing *Bcl11b*, *Cntn6*, and *Foxo1* (Piri1); a cluster of cells spatially concentrated in cortical layers V and expressing *Satb2* (Exc5); a cluster of cells spatially concentrated in cortical layers V and VI and expressing *Tle4*, *Bcl11b*, and *Fezf2* (Exc-5,6); a cluster of cells spatially concentrated in cortical layer VIb and piriform layers 2 and 3, and expressing *Nr4a2* and *Ccn2* (Exc-6b,P2,3); a cluster of cells spatially located in cortical layers 6b and the subplate, and expressing *Tle4*, *Nr4a2*, *Satb2, Rgs8,* and *Ccn2* (Exc-6b,SP); and a cluster of cells spatially located in the subplate and expressing *Nr4a2*, *Satb2* (SP; Figures 2C, 2F, and S3B). Interneurons formed three subtype clusters, including unique populations with high expression of *Pvalb* (Int1), *Sst* (Int2), or *Prox1* (Int3) (Figure 2C, 2E and S3A).

OL-lineage clusters from the cortex and corpus callosum were subclassified as OPCs by the expression of *Pdgfra* and *Sox10*, and OLs by the expression of *Mag* and absence of *Pdgfra* (Figure 2B). OL clusters corresponding to different stages of OL-lineage progression had graded expression patterns for many subtype-specific genes. Clusters with the highest enrichment for *Fyn* and *Gpr17* were classified as newly formed OLs, clusters enriched for *Tcf7l2* and *Myrf* were classified as immature OLs, and clusters enriched for *Trf*, *Car2*, and *Opalin* were classified as mature OLs. In the cortex, we distinguished a cluster of late OPCs (pre-OLs) expressing low levels of the OPC marker *Pdgfra* and newly formed OL markers *Fyn* and *Gpr17* (Figure 2D and S3C).

Astrocyte clusters in the cortex and corpus callosum were subclassified as reactive astrocytes if they expressed *Serpina3a, C3, Ggta1*, and/or high *Gfap*. In the cortex, reactive astrocytes formed two clusters, with one population (Astro-react1) expressing *Tnc*, *Vcam1*, *Sox9*, and *Notch1/2*, and a second population (Astro-react2) enriched for *Gfap* and *Sparc* (Figure 2D and S3D). In the corpus callosum, we identified one cluster of reactive astrocytes (Astro-react) which expressed *Vcam1*, *Sox9*, and *Sox2*, resembling Astro-react1 of the cortex (Figure 3F and S3K). Astrocytes that did not express reactive astrocyte markers were classified as homeostatic astrocytes. In the cortex, we identified three clusters of homeostatic astrocytes: one population (Astro-hom1) was enriched for *Slc1a3*, one population (Astro-hom2) was enriched for *Cx3cl1*, and one population (Astro-hom3) was enriched for *Sparc* (Figure 2D and S3D). In the corpus callosum, we identified three clusters of homeostatic astrocytes: one population (Astro-hom1) was enriched for *Slc1a3*, one population (Astro-hom2) was enriched for *Clu*, *Ncan* and *Bcan*, and one population (Astro-hom3) was enriched for *Sparc* (Figure 3F and S3I).

**Figure 3:**
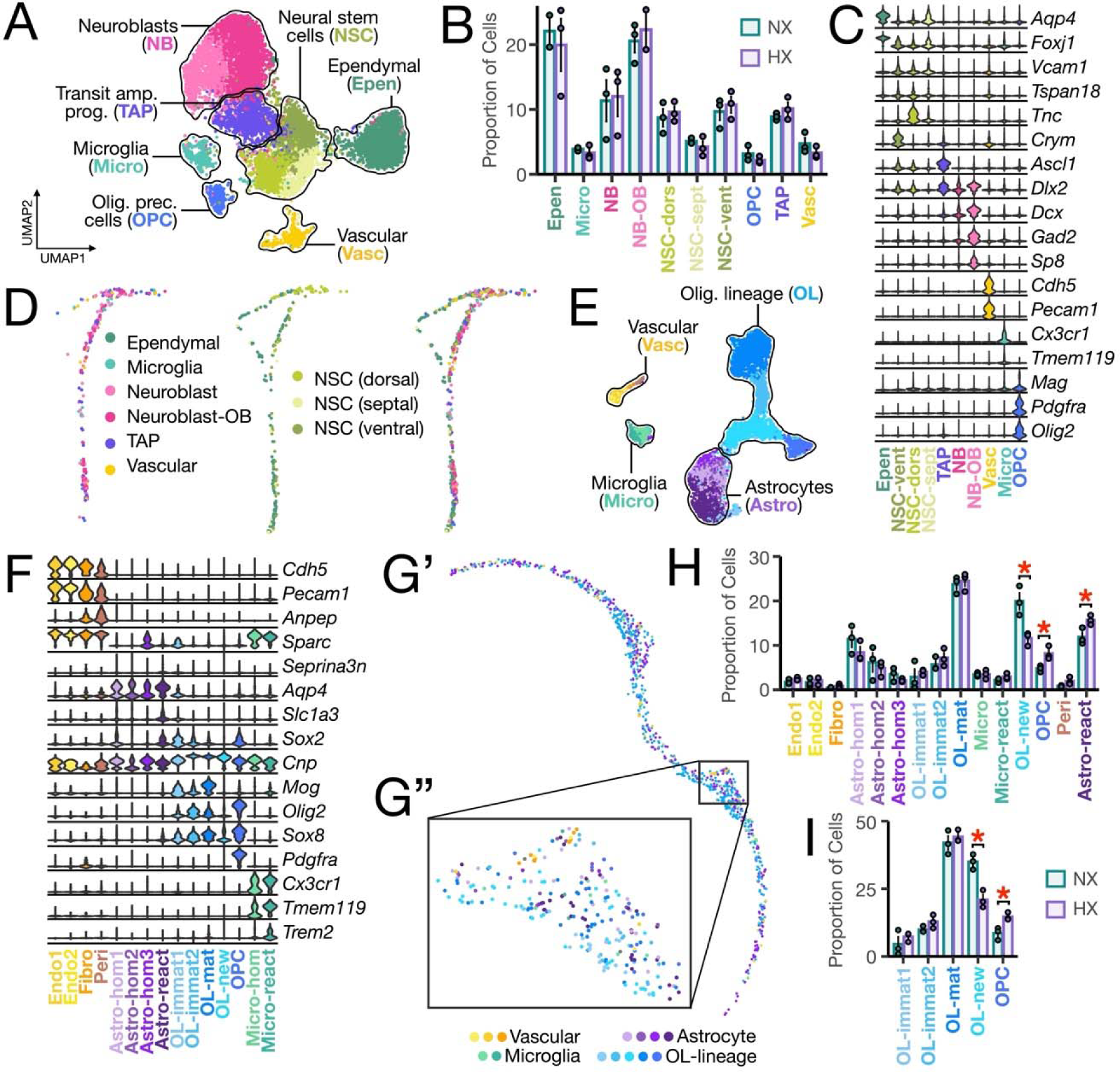
Cell type classification and spatial distribution in the P21 mouse SVZ and corpus callosum reveals arrested OL differentiation in the corpus callosum following perinatal hypoxia. **(A)** UMAP visualization of all SVZ cells, for a total of 10,754 cells. UMAP plot is generated from combined replicates across NX and HX conditions. Each cluster is colored by cell type and are labelled by cell type, with all NSCs labelled together and all neuroblasts labelled together, as follows: Epen: ependymal cell, TAP: transit amplifying progenitor, NB: neuroblast, NB-OB: olfactory bulb neuroblast, NSC: neural stem cell, NSC (dors): dorsal neural stem cell, NSC (sept): septal neural stem cell, NSC (vent): ventral neural stem cell. **(B)** Bar plot displaying the proportion of Cell Subtypes in the SVZ of HX versus NX mice. The x-axis displays all major Cell Subtypes profiled through MERFISH. Data obtained from *n*L=L3 male biological replicates per condition. Bars represent the average values for each condition, and dots represent the average values for each mouse per condition. Error bars represent the average ± 1 standard deviation. Each bar is labeled by cell subtype as previously defined. **(C)** Violin plot showing gene expression patterns of Cell Subtypes identified in the SVZ with several key canonical marker genes used for cluster identification. The width of the violin corresponds to the proportion of cells expressing the indicated gene, and the color of the violin corresponds to the cell subtype. Each column is labeled by Cell Subtype as previously defined. **(D)** Spatial plot displaying the distribution of Cell Subtypes in the SVZ. **(E)** as in (A) but displaying all corpus callosum cells, for a total of 8,708 cells. Each cluster is colored by Cell Type as follows: Astro: astrocyte, Micro: microglia, OL: oligodendrocyte, Vasc: vascular. **(F)** as in (C) but displaying gene expression patterns of Cell Subtypes identified in the corpus callosum. **(G’)** Spatial plot displaying the distribution of Cell Types in the CC. **(G”)** Zoomed view of the panel delineated in G’. Bar plots displaying the proportion of **(H)** Cell Subtypes **(I)** OL-lineage Cell Subtypes of HX versus NX mice.

Microglia in the cortex and corpus callosum were subclassified as either reactive if they expressed reactive microglia markers *Trem2* and *Ptprc*, or homeostatic if they lacked those markers. Corpus callosum microglia formed one homeostatic (Micro-hom) and one reactive cluster (Micro-react) (Figure 3F and S3K), whereas cortical microglia formed one homeostatic cluster (Micro-hom) and two reactive clusters: one cluster of reactive microglia was enriched for *Sirpa* (Micro-react1), and the other (Micro-react2) was enriched for *Sox9* and neuron- (*Grin1*, *Gria2*) and OL-specific (*Cnp*, *Plp1*) genes (Figure 2D, S4E). Neuron- and OL-specific mRNA in Micro-react2 cells suggest possible OL and neuron engulfment. Conversely, because *Sirpa* expression in microglia inhibits phagocytosis^23,24^, Micro-react1 may represent a less phagocytic subtype of microglia.

Vascular cell clusters in the cortex and corpus callosum were subclassified as pericytes (Peri) if they expressed *Pdgfrb* and *Atp13a5*, fibroblasts (Fibro) if they expressed *Col1a1*, *Col1a2*, *Fbln1*, or *Fbln2*, and endothelial cells if they expressed high levels of *Pecam1*, *Esam*, and *Cdh5* but none of the pericyte or fibroblast markers. In the cortex, fibroblasts formed multiple clusters including a cluster that was enriched for *Fbln1*, *Gfap*, and *Pdgfra*, low in *Pecam1* and *Esam*, and was spatially enriched in the meninges (Fibro-1); a cluster that was enriched for *Icam1* and *Gfap* (Fibro-2); a cluster that was enriched for *Ccn2*, *Cspg4*, and *Tgfb3* (Fibro-3); and a cluster that was enriched for *Fbln2*, *Vcam1*, and *Edn1* (Fibro-4). Cortical endothelial cells also formed multiple clusters including a cluster that was enriched for *Cxcl12* and *Cx3cl1* (Endo-1), a cluster enriched for *Flt1* (Endo-2), and a cluster enriched for *Cd82* and *Serpine1* (Endo-3; Figure 2D, S4E).

In the SVZ, we identified vascular cells, microglia, ependymal cells, quiescent neural stem cells (NSCs), transit amplifying progenitors (TAPs), OPCs, and both olfactory bulb (OB) and non-OB neuroblasts (NB-OB, Figure 3A). Quiescent NSCs were identified by their expression of *Vcam1*, *Thbs4*, *Tspan18*, and *Slc1a3* and absence of *Mki67*, transit amplifying progenitors (TAPs) by their expression of *Mki67*, *Egfr*, *Ascl1* and *Dlx2*, neuroblasts by their expression of *Dlx2* and *Dcx* and absence of *Ascl1*, and ependymal cells were identified by their expression of *Foxj1* (Figure 3C). NSCs were analyzed independently and formed 3 clusters: one cluster expressing *Crym* and spatially localized to the ventrolateral wall of the SVZ (NSC-vent), a cluster enriched in *Tnc* and spatially localized to the dorsal wall of the ventricle (NSC-dors), and the third cluster was enriched for *Ednrb* and *Aqp4* and spatially localized to the septal wall of the ventricle (NSC-sept) (Figure 3D and S3G). NB-OB were identified based on the expression of *Sp8* whereas non-OB neuroblasts lacked *Sp8* expression (Figure 3C). The latter group may represent neuroblasts that will migrate to the caudoputamen.^25,26^

### Region-specific patterns of signaling-related gene expression

Recent studies have suggested that cellular phenotypes are at least partly informed by spatial location in the brain.^13,19,20^ Before investigating how HX alters region-specific cellular signaling profiles, we first compared transcriptional profiles of non-neuronal cells between distinct brain regions (SVZ, corpus callosum, superficial layers of the cortex, and deep layers of the cortex) in the P21 NX brain to identify baseline regional heterogeneity. This was done by differential gene expression analysis where a false discovery rate (FDR)< 0.05 and log_2_ fold-change (log_2_(FC)) > 0.25 were used to identify statistically significant differences in each comparison (see *Methods*). We observed extensive differences in signaling-related gene expression levels across all regions and cell types. Notably, vascular cells were remarkably different between the upper and deep cortex with respect to expression of genes involved in WNT, Insulin-like growth factor (IGF), BMP, and fibroblast growth factor (FGF) signaling. Upper cortex vascular cells were enriched for expression of *Fzd1/2/7*, *Wnt4/5a*, *Sfrp1*, *Igf2*, *Igfbp2*, *Bmp4/5/7*, *Fgf1*, *Fgfr1*, and *Fgfr2*, while deep cortex vascular cells were enriched for expression of *Igf1r* and *Apc* (Figure S4F and Table S2). Compared to all other regions, vascular cells of the upper cortex but not the deep cortex were enriched for WNT ligand (*Wnt5a*), Bmp ligand (*Bmp4/5/7*), and IGF ligand (*Igf2*, *Igfbp2*) genes (Figure S4 and Table S2). These results suggest activation of the canonical WNT pathway within vascular cells in the upper cortex, and robust WNT inhibition mediated by the negative regulator *Apc*, which was expressed in vascular cells in the deep cortex.

OPCs in different regions exhibited unique patterns of signaling-related gene expression. For example, SVZ and corpus callosum OPCs expressed higher levels of *Bmp4* and *Notch1*, while cortical OPCs expressed higher levels of *Cx3cl1* (Table S2). Microglia also showed notable differences between regions. SVZ microglia exhibited higher expression of genes involved in WNT signaling, such as *Fzd1/2/8*, *Lrp4/6/8*, *Sfrp1*; genes involved in sonic hedgehog (SHH)signaling such as *Ptch1*, *Gli1/3*; and Ephrin signaling genes such as *Efna2*, *Efnb2/3*, and *Epha4* (Figure S4A-C and Table S2). This may suggest that in the developing mouse brain, microglia in the SVZ are uniquely suited for sensing and regulating the SVZ niche to modulate neurogenesis.^27^

### HX-associated changes in cellular composition

We next assessed differences in global cell type proportions between HX and NX by leveraging the multiple biological replicates for each condition (Figure 3, see *Methods*). In the corpus callosum, we found that the proportion of newly formed OLs was markedly reduced following HX (50% reduction, p = 0.016), while the proportion of OPCs was significantly increased (40% increase, p = 0.025; Figure 3H and 3I). These findings are consistent with previous literature suggesting that chronic sublethal neonatal HX induces WMI followed by a robust period of oligodendrogenesis.^7,9^ We also observed a significant increase in the proportion of reactive astrocytes in the corpus callosum in HX (Figure 3H). This is consistent with previous work showing that there is an increase in the number of *C3*-expressing reactive astrocytes in the corpus callosum following acute perinatal WMI.^28,29^

In the cortex of HX mice, we identified a significant increase in the abundance in Endo-1 cells, the most abundant subpopulation of endothelial cells (82% increase, p = 0.015, Figure 2I), suggestive of vascular remodeling. Lastly, we observed a significant increase in the proportion of a large population of excitatory neurons in layer II-III (18% increase, p = 0.019), and a significant decrease in a small population of excitatory neurons in layers I-III (54% reduction, p = 0.049, Figure 2K), indicating a change in the cell type composition of upper layer excitatory neurons with an overall increase in excitatory neurons. We also observed an increase in the proportion of Int1, a population of cortical interneurons (11% increase, p = 0.038, Figure 2K). This could suggest a disruption or delay in programmed neuron death and clearing in the upper cortical layers (the last layers to form and mature), a process that largely takes place in the first week after birth, coinciding with the timing of HX exposure.^30,31^

### Proximity analysis reveals widespread changes in local cellular architecture

We next evaluated changes in local cellular neighborhood composition associated with perinatal HX. Unlike spot-based spatial transcriptomics methods, MERFISH enables us to accurately map both spatial locations and transcriptional identities of individual brain cells. We leveraged this information using a novel computational approach for quantifying condition-associated differences in local cell type proximities (see *Methods*), which addresses some limitations of existing methods.^32,33^ Briefly, we calculated the condition-related differences in the proportion of cells of one type in the vicinity of a second type, thus capturing asymmetrical differences in local proportions between cell types. For example, our approach can detect the depletion of astrocytes surrounding neurons when the reverse is not true. Second, we directly compare cell type proximities between conditions using a general linear mixed effects model, accounting for the statistical non-independence of measures from the same tissue sample when there are multiple biological replicates (see *Methods*). We applied this approach at the microscale, capturing local changes in proximity that matter for cellular interaction, however, we note that such differences may also reflect global cellular abundance differences between conditions (e.g., those shown in Figure 2 and 3).

In the cortex, perinatal HX was associated with decreased proximity of homeostatic astrocytes to several cell types (Figure 4A), including pre-OLs (beta = -0.46, FDR = 8.45 ⍰ 10^-5^; Figure 4D), newly formed OLs (beta = -0.48; FDR = 0.012), and immature OLs (beta = -0.43, FDR = 4.21 ⍰ 10^-3^). This may indicate a disruption in the supportive interactions provided by astrocytes, such as the secretion of local factors (e.g. BDNF, CNTF) that promote OL maturation and myelination during healthy conditions and following injury[REF].^34^ HX induced dramatic changes in the proximity of different populations of cortical excitatory neurons to other neurons and non-neuronal cells. For example, after HX, a population of upper layer excitatory neurons, Exc-1,2,3, had decreased proximity to interneurons (beta = -0.15, FDR = 6.65 ⍰ 10^-6^), reactive (beta = -0.06, FDR = 0.046) and homeostatic astrocytes (beta = - 0.13, FDR = 1.07 ⍰ 10^-3^), and pre-OLs (beta = -0.05, FDR = 1.93 ⍰ 10^-3^).

**Figure 4:**
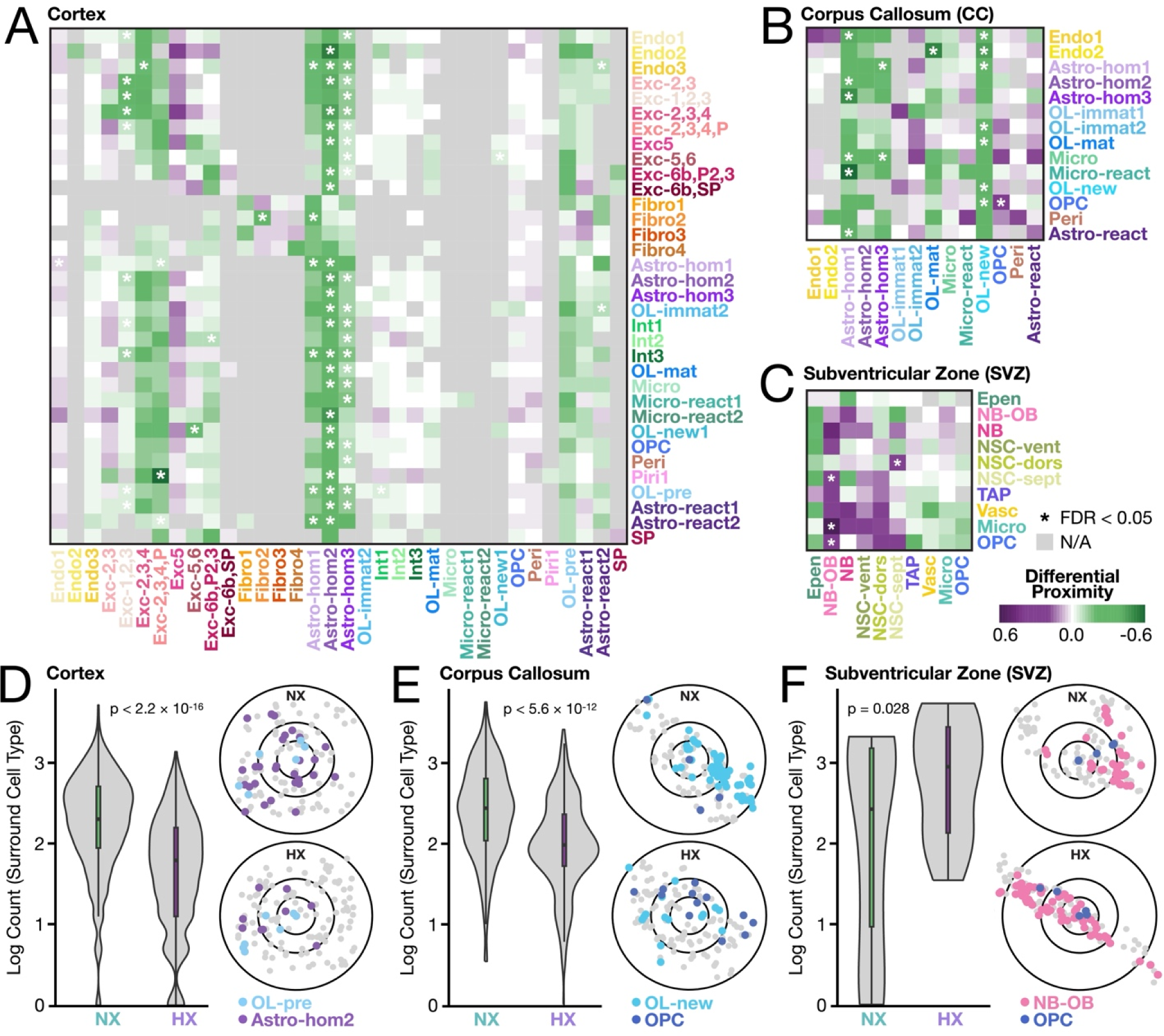
Perinatal hypoxia alters local spatial proximity between select Cell Type pairs across brain regions. Differential proximities between cell type pairs (heatmaps) in the **(A)** cortex, **(B)** corpus callosum, and **(C)** SVZ. For cortex and corpus callosum, Cell SubType classification was used. Positive scores (purple) indicate an increased proportion of the surround cell type (columns, x-axis) in proximity of the center cell type (rows, y-axis) in HX relative to NX, whereas negative scores (green) indicate a decrease. Grey indicates not enough non-zero data was present for the respective cell type pair. Significant (FDR < 0.05) relationships are denoted with an asterisk. Exemplar spatial plots (right panels) indicating the distribution of select center and surround cell types between NX (top) and HX (bottom) in **(D)** cortex, **(E)** corpus callosum, and **(F)** SVZ. The outer circles in the spatial plots correspond to the radius used to calculate the differential proximities in (A-C) and have been normalized to reflect the median cellular distances per sample. Violin plots (left panels) show the counts of the surround cell type in proximity to every central cell across the sample corresponding to the spatial plot (not just the sample area shown). P-values for t-tests are shown.

In the corpus callosum following HX (Figure 4B), we observed significantly reduced proximity of newly formed OLs and homeostatic astrocytes in the neighborhoods of several cell types, including OPCs (beta = -0.37, FDR = 6.82 ⍰ 10^-3^; Figure 4E) and endothelial cells (beta = -0.65, FDR = 0.011), as well as reactive astrocytes (beta = -0.57, FDR = 2.62 ⍰ 10^-13^) and microglia (beta = -0.45, FDR = 0.016). Such local interactions are likely essential for OL maturation and efficient myelination.^34^ The altered spatial organization of these cells potentially contributes to the impaired myelination characteristic of diffuse WMI.

In the post-HX SVZ, NSCs showed significantly increased proximity to differentiating neuroblasts (beta = 0.28, FDR = 7.12 ⍰ 10^-3^), with a trending increase in proximity to OPCs, microglia, and vasculature (Figure 4C and 4F). These changes align with the roles of activated microglia in promoting oligodendrogenesis and neurogenesis. The closer association with vascular components may indicate an adaptive response to HX, possibly involving angiogenesis or enhanced metabolic support. Neuroblasts saw increased proximity to TAPs (beta = 0.36, FDR = 1.61 ⍰ 10^-4^), microglia (beta = 0.81, FDR = 6.90 ⍰ 10^-4^), and OPCs (beta = 0.54, FDR = 0.026). These changes may reflect neural repair through enhanced interactions of neuroblasts with TAPs for proliferation, with microglia for inflammation resolution, and with OPCs for myelination, altogether accelerating recovery processes. Overall, these observations suggest a reorganization or adaptation of the SVZ niche in response to perinatal HX, highlighting the region’s plasticity under developmental stress.

In total, our proximity analysis emphasizes a crucial spatial component to the effects of perinatal HX beyond gene expression. Changes were extensive and heterogeneous across cell types and regions, ranging from disrupted myelination in the corpus callosum to adaptive cellular reorganization in the SVZ.

### Perinatal HX alters regional patterns of signaling-related gene expression

We next analyzed cell type- and region-specific transcriptional changes between HX and NX conditions using differential gene expression analysis (see *Methods*). This comparative analysis aimed to identify molecular processes perturbed during HX exposure in the context of early brain development. In all cell type populations assayed in the SVZ, corpus callosum, and cortex, we found that the vast majority of differentially expressed genes (DEGs) were upregulated (i.e., had increased expression) following HX (Figure 5A-F, Table S2).

**Figure 5:**
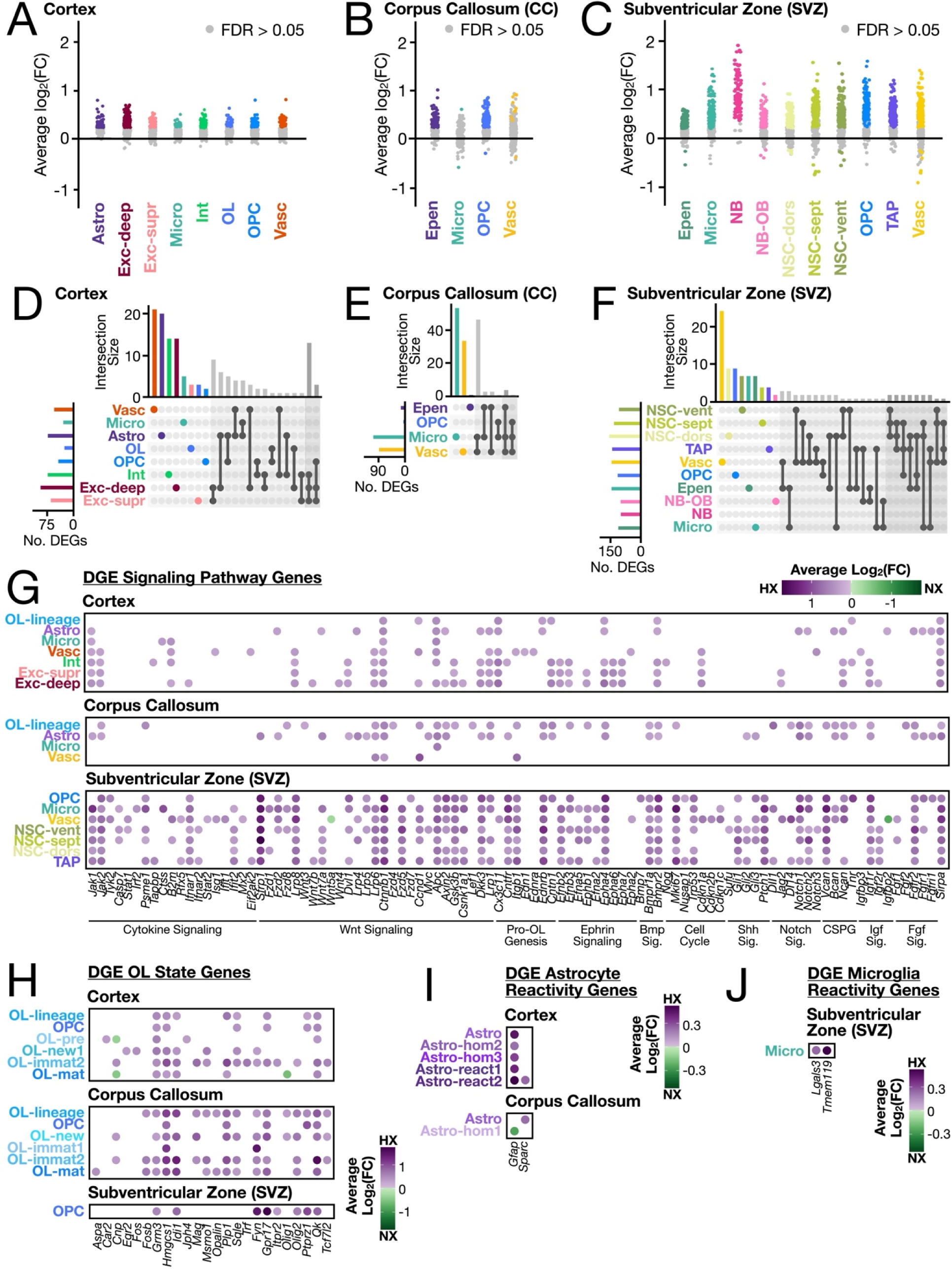
Differential gene expression analysis finds transcriptionally distinct cellular subpopulations in the HX brain. **(A-C)** Strip plots displaying DEGs between P21 HX and NX mice. Each dot represents a DEG: colored dots represent significant genes (FDRL<L0.05) per cell type, gray dots represent non-significant genes (FDR > 0.05). The x-axis displays all major cell types across the **(A)** cortex, **(B)** corpus callosum, and **(C)** SVZ regions profiled through MERFISH analysis. Data obtained from *n*L=L3 biological replicates per condition. **(D-F)** UpSet plots displaying the number of unique and shared DEGs across cell types in the **(D)** cortex, **(E)** corpus callosum, and **(F)** SVZ. Unique genes are colored based on cell type, and genes shared between two or three cell types indicated by black dots connected by lines according to shared origins. The histogram indicates the number of DEGs for each cell type, and the barplots show the number of significant DEGs (FDRL<L0.05) per cell type. Differential gene expression comparison of HX and NX in each region (cortex, corpus callosum, and SVZ) for (**G)** signaling genes across Cell Types, **(H)** OL state genes across all OL-lineage cells (from Cell Type) and Cell Subtypes within the OL-lineage **(I)** astrocyte reactivity genes across all astrocytes (Astro from Cell Type) and Cell Subtypes within astrocytes, and **(J)** microglia reactivity genes across all microglia (Micro from Cell Type) and Cell Subtypes within astrocytes. Genes in **(H)** are grouped and labeled according to the signaling pathway or function that they participate in. Cortex and corpus callosum data in (A, B, D, E, G) display Cell Type information, and in (H, I) display both Cell Type and Cell SubType information. Only significant genes are shown. Colors represent average log_2_(FC) as per legend. Results are also shown in Table S2.

We expanded our findings from the differential gene expression analysis by focusing on understanding cell-cell signaling after perinatal HX. Neighboring cells communicate via secreted molecules (ligands) that can act on receptors in nearby cells. These ligand-receptor (LR) networks are fundamental signaling nodes that regulate circuit function and injury response. To understand how local cell-cell communication networks change in the context of perinatal brain injury, we performed a spatially informed LR analysis using CellChat v2 (see *Methods*). We grouped predicted LR pairs based on whether signaling occurred within the same cell type (e.g., OPC to OPC; intra-lineage) or between different cell types (e.g., microglia to OPC; inter-lineage). After exposure to perinatal HX, we identified region-specific cellular programs governing regenerative oligodendrogenesis and neurogenesis.

In the SVZ, we found that HX induced an upregulation of *Egfr*, *Lmnb1*, and the proliferation marker *Mki67* within most SVZ cell types, including NSCs, OPCs, and TAPs, suggestive of NSC activation (Figure 5G, Table S2).^35^ In response to HX, all SVZ neural progenitors, including NSCs, OPCs, and TAPs, upregulated genes involved in OL differentiation such as *Qk* and *Ptprz1*, which suggests priming towards an OL fate. Neural progenitors also showed increased expression of genes known to be expressed in early neuron development including *Dcx*, *Gria2*, and *Stmn1*, suggestive of increased neurogenesis. We observed dramatic changes in BMP, WNT, and Notch signaling, consistent with their role in regulating stem and progenitor cell maintenance, proliferation, differentiation, and fate decisions (Figures 5G and 6E-F, Tables S2 and S3). Specifically, there was a significant increase in *Bmp7* expression in OPCs and an increase in BMP7 and BMP4 signaling from OPCs to OPCs, NSCs, TAPs, microglia, and vascular cells (Figures 5G and 6E-F). In addition, the WNT ligand *Wnt7a*, the receptor *Lrp6*, the effector *Ctnnb1*, and target gene *Ccnd1* all demonstrated increased expression in most SVZ cells, including NSCs, after HX which translated to a widespread increase in WNT signaling (Figure 6F, Table S3). In the HX SVZ, *Notch1* and *Notch2* receptors were upregulated in most cell types, while the ligand genes *Dll1* or *Dll4* were upregulated in vascular cells, TAPs, and immune cells (Figure 5G, Table S2) which translated to an increase in Notch signaling between these cell types (Figure 6F, Table S3). Interestingly, we found increased transforming growth factor β 2 and 3 (TGFβ-2/3) signaling between the vasculature and microglia in the SVZ after HX (Figure 6F, Table S3). TGFβ is a critical component of the microglial pro-neurogenic response.^36–38^

**Figure 6:**
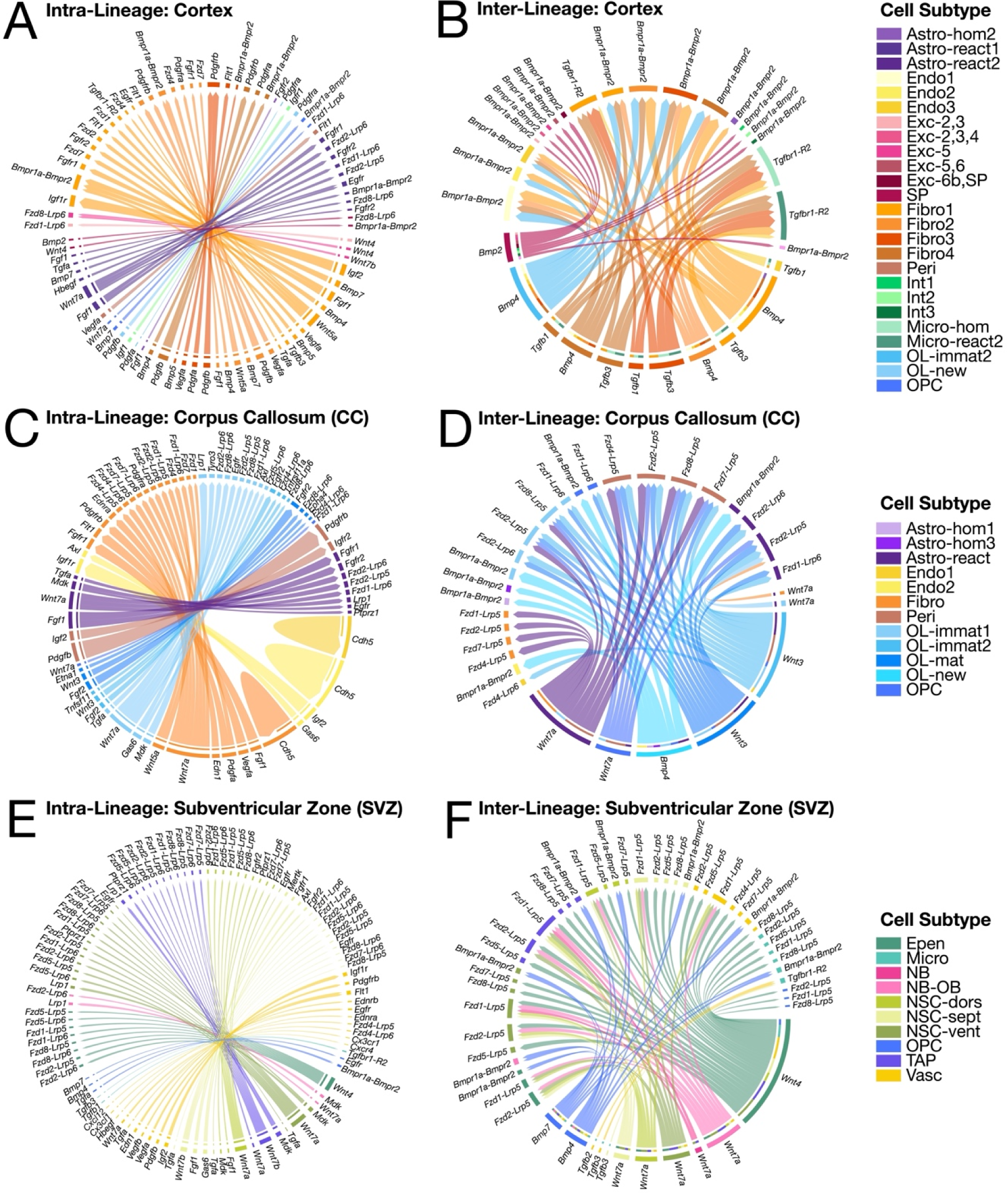
Cell subtypes exhibit discrete cell-cell signaling networks in the brain of mice exposed to postnatal hypoxia. Chord diagrams of predicted intra-lineage cell-cell communication pathways that are upregulated in the **(A)** cortex, **(B)** corpus callosum, and **(C)** SVZ of HX versus NX mice. Chord diagrams of predicted inter-lineage cell-cell communication pathways that are upregulated in the **(D)** cortex, **(E)** corpus callosum, and **(F)** SVZ of HX versus NX mice. Cell Subtype identity of the ligand is indicated in the outermost edge of the diagram, while the Cell SubType identity of the receptor is indicated by the internal ring. Encoding gene symbols are used to represent predicted interactions of their protein products. Colored arrows indicate the specific ligand-receptor (LR) pairs and are colored according to the outgoing ligand signal. For all chord diagrams, only significant ligand and receptor interactions are plotted (p < 0.05). Full results are shown in Table S3.

In the corpus callosum, we identified several signaling nodes that promote white matter recovery after perinatal HX, including upregulation of MAG to MAG and TGF-α to epidermal growth factor receptor (EGFR) signaling between Cell SubTypes of OL-lineage cells (Figure 6D, Table S3).We observed increased WNT signaling from astrocytes and OLs, increased IGF2/IGF1R signaling from vascular cells to astrocytes and OLs, and increased CNTN1*-*Notch signaling between OLs, astrocytes, and vascular cells. Changes in Notch signaling was a commonly identified theme: *Notch1* and *Notch2* were upregulated in astrocytes and OL-lineage cells and the ligand *Dll1* was upregulated in OL lineage cells (Figure 5G, Table S2). Moreover, we found increased GAS6 signaling through its receptors (TAM receptors, encoded by *Tyro3*, *Axl*, *Mer*) between microglia and astrocytes, vascular cells, and OL-lineage cells (Table S3). Following demyelinating injury, GAS6-TAM signaling promotes remyelination and glial cell development, including suppression of a deleterious (anti-regenerative) microglia response.^39,40^ Notably, we found evidence of neurovascular adaptation after HX, including upregulation of genes involved in cell migration and vascular permeability (*Fstl1* and *Itgb1*)^41^ in vascular cells in the corpus callosum, which may promote immune cell infiltration and/or guide the migration of new neurons and glial cells (Figure 5G, Table S2). We observed an increase in inflammatory and migratory signaling from corpus callosum fibroblasts, including upregulation of Interleukin 34 (IL-34)to colony stimulating factor 1 receptor (CSF1R) and ephrin A1 (EFNA1) to Ephrin type-A receptor 7 (EphA7)ligand-receptor pairs, both of which may support glial scar formation (Figure 6C-D, Table S3).^42–44^

In the cortex, LR analysis was notable for a broad, multicellular increase in several signaling pathways previously implicated in brain circuit wiring and the response to injury. Endothelin-1 (EDN1)/Endothelin Receptor Type B (EDNRB) signaling was increased between endothelial cells, fibroblasts, astrocytes, OLs, and microglia, and astrocytes (Figure 6A-B, Table S3). Endothelin regulates myelination capacity in OLs, as endothelin is upregulated after demyelination and directly inhibits OPC differentiation via astrocytes.^45^ Additionally, we found that fractalkine (encoded by *Cx3cl1*)-CX3CR1 signaling, a regulator of gliogenesis, was increased between several non-neuronal cell populations, including OLs, OPCs, fibroblasts, and astrocytes, to reactive microglia (Figure 6B, Table S3). A regulator of fractalkine expression and remyelination, TGFβ1 signaling, was also enhanced following HX between endothelial cells, fibroblasts, and microglia. Canonical WNT signaling is important for TGFβ-mediated fibrosis^46^, and we identified increased WNT signaling in deep layer cortical excitatory neurons (WNT4 with FZD4), as well as between vascular cells, deep layer excitatory neurons and astrocytes (WNT7A with FZD4). We observed a HX-induced increase in cytokine signaling (including IGF2-IGF1R, IGF2-IG2R, fractalkine-CX3CR1, TGFB1-TGFBR1, TGFB3-TGFBR1 ligand-receptor pairs) largely sourced by fibroblasts and other vascular cells, and targeting vascular cells and microglia (Figure 6A-B, Table S3). Thus, vascular cells may represent an important source of inflammatory signaling leading to increased microglial activation after perinatal HX. Additionally, several homophilic (CDH5-CDH5, ESAM-ESAM, PECAM1-PECAM1) and heterophilic (FGF1/2-FGFR1/2R, BMP4/7-BMPR1A, VEGFA-FLT1) endothelial interactions involved in regulating vascular development and blood brain barrier permeability integrity were dysregulated in all three regions of the brain (Figure 6, Table S3).^47–50^ In addition, in the cortex, we found that HX induced an upregulation of *Sst* expression in interneurons and an upregulation of *Sstr2* expression in both interneurons and excitatory neurons (deep and superficial layers). Furthermore, LR analysis revealed increased *SST*-*SSTR2* signaling from interneurons to other neurons following HX (Table S3), all of which could impact excitatory-inhibitory neuron balance.

Ephrin communication was commonly altered across the cortex, SVZ, and corpus callosum. Cortical excitatory neurons and interneurons in the HX brain showed an upregulation of receptors *Epha4*, *Epha6*, and *Epha7* and ligands *Efnb2* and *Efnb3* (Figure 5G, Table S2). EFNA1 signaling by cortical fibroblasts to OL-lineage cells and astrocytes was significantly upregulated post-HX (Table S3). In the SVZ, there was widespread upregulation of ephrin genes; for example, *Epha4* and *Efnb2* were upregulated in all cell types following HX. Furthermore, all SVZ cells participated in ephrin signaling as sources and targets following HX. The OL-lineage in the corpus callosum also upregulated ephrin genes and showed increased ephrin signaling targeting all major Cell Types (Table S3). EFNA1 interacts with EphA4 to inhibit OL process extension via ephexin1-RhoA-Rock-myosin 2.^51^ These HX-induced ephrin interactions may play an important role in guiding the migration and process extension of injury-responsive glial cells or new neurons.

Another important source of cell guidance signaling is the extracellular matrix. In particular, chondroitin sulfate proteoglycans (CSPGs) are extracellular matrix components known regulate cell migration, synapse formation and maturation, and inflammation in the developing brain.^52,53^ However, the upregulation of CSPGs following brain injury has been shown to block migration of neuroblasts and inhibit regeneration.^52,54^ We found that OL-lineage cells and astrocytes in the cortex and corpus callosum, cortical excitatory neurons, cortical interneurons, and SVZ OPCs showed an upregulation of CSPG genes such as *Vcan*, *Ncan*, *Bcan*, and *Tnr* (Figure 5G, Table S2). As such, there may be a change in neural guidance cues that could affect the appropriate migration and integration of new cells in the HX brain.

### Perinatal HX is associated with changes in region-specific cell states

Finally, we asked whether microglia, astrocyte, and OL-lineage cell state is dictated by spatial context by comparing gene expression patterns between the SVZ, corpus callosum, and cortex. We found notable differences in cell state associated gene expression across different anatomical regions in the NX brain. Differential gene expression analysis revealed that microglia in the NX SVZ and corpus callosum exhibited a significantly lower expression of both *Tmem119* and *Trem2* compared to microglia in the superficial cortex and deep cortex (Figure 7Q-T). SVZ microglia showed the same trend of lower expression of both *Tmem119* and *Trem2* when compared to the corpus callosum (Figure 7Q). This trend is unexpected because it is widely acknowledged that *Tmem119* has higher expression in homeostatic microglia while *Trem2* is higher in reactive microglia.^55^ However, recent work showing that *Tmem119* gene expression levels and spatial distribution in the brain are incongruent with protein levels following traumatic brain injury challenge the use of transcriptomic *Tmem119* as a marker for homeostatic microglia.^56^ Therefore, to better characterize the reactivity state of microglia across different regions, we calculated reactivity scores using multiple genes shown previously to be specific to or enriched in reactive microglia at the transcriptomic level (see *Methods*).^57,58^ We observed a lower reactivity score in corpus callosum microglia, specifically Micro-react, compared to superficial cortex microglia (Figure 7U-W). HX exposure reduced the reactivity score of corpus callosum microglia significantly, pushing microglia reactivity in the corpus callosum to be significantly lower than both upper and deep cortex microglia. In addition, only microglia in the SVZ exhibited a significant change in state related genes following HX exposure, including an upregulation of homeostatic microglia-associated *Tmem119* and reactive marker *Lgals3* (Figure S4C).

**Figure 7:**
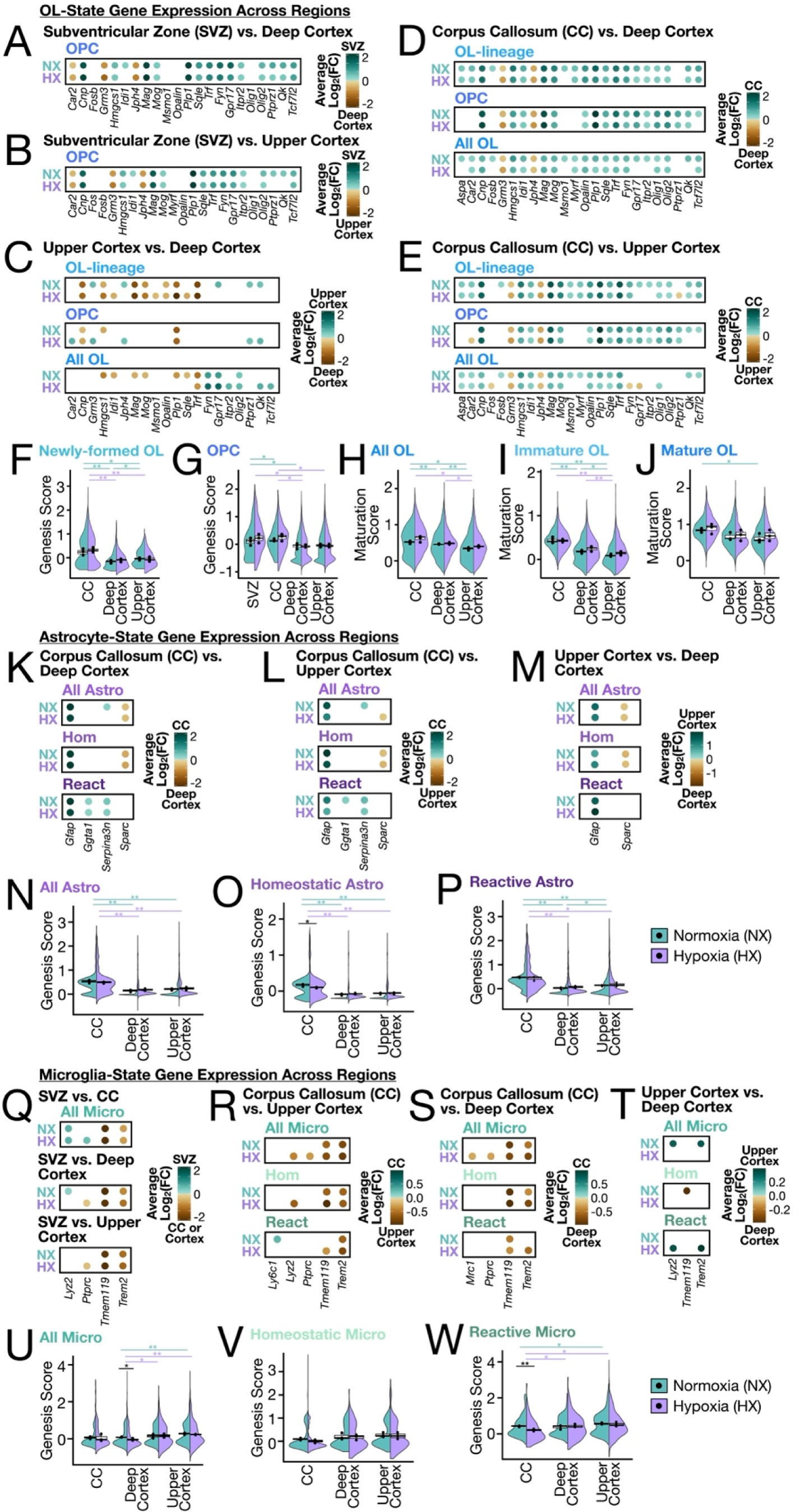
Cell state changes of OL-lineage cells, astrocytes, and microglia across anatomical regions and following perinatal hypoxia exposure. Dotplots displaying differential gene expression results of OL-state (OL genesis and OL maturation) genes comparing all OL-lineage cells (Cell Type) and Cell Subtypes within NX and HX between **(A)** SVZ and deep cortex, **(B)** SVZ and upper cortex, **(C)** upper and deep cortex, **(D)** corpus callosum and deep cortex, and **(E)** corpus callosum and upper cortex. Violin-boxplots showing OL genesis state score in **(F)** newly-formed OLs and **(G)** OPCs, and OL maturation score of **(H)** all OLs, **(I)** immature OLs, and **(J)** mature OLs, split by condition across the SVZ, corpus callosum, and cortex. Dotplots displaying differential gene expression results of astrocyte reactivity genes comparing all astrocyte Cell Types and Cell Subtypes within NX and HX between **(K)** corpus callosum and deep cortex, **(L)** corpus callosum and upper cortex, **(M)** upper and deep cortex. Violin-boxplots showing astrocyte reactivity score of **(N)** all astrocytes, **(O)** homeostatic astrocytes, and **(P)** reactive astrocytes, split by condition, across the SVZ, corpus callosum, and cortex dotplots displaying differential gene expression results of microglia reactivity genes comparing all microglia Cell Types and Cell Subtypes within NX and HX between **(Q)** SVZ and corpus callosum, SVZ and deep cortex, SVZ and upper cortex, **(R)** corpus callosum and deep cortex, **(S)** corpus callosum and upper cortex, **(T)** upper cortex and deep cortex. Violin-boxplots showing microglia reactivity score of **(U)** all microglia, **(V)** homeostatic microglia, and **(W)** reactive microglia, split by condition, across the SVZ, corpus callosum, and cortex. For all dotplots, only significant genes are shown (FDR < 0.05). Dot color represents average log_2_(FC) as per the associated legend. For all violin-boxplots, the width of the violin plot represents the proportion of cells expressing that value of the score. Each dot represents the average score for each replicate, the bold central line of the boxplot represents the median score for the condition, and the limits of the box represent the interquartile range. Significance is determined using the two-tailed Student’s t-test. *p-value < 0.05; **p-value < 0.01. Results for all dotplots are also shown in Table S2.

In the NX brain, all astrocytes in the corpus callosum had significantly higher expression of *Gfap* than astrocytes in the cortex, with Astro-react also exhibiting higher levels of reactive markers *Ggta1* and *Serpina3n* in the corpus callosum (Figure 7K-M). We then calculated astrocyte reactivity scores from established reactive astrocyte genes^59–61^ and found that astrocytes in the NX corpus callosum exhibited a significantly higher state of reactivity compared to those in the cortex. This heightened reactivity was not altered by HX treatment. Within the cortex, astrocytes in the superficial layers showed a slightly elevated reactivity score over those in the deep layers (Figure 7N-P). Astro-hom, on the other hand, had higher *Sparc* expression in the cortex than in the corpus callosum, with deep layer astrocytes expressing more *Sparc* than superficial ones, a trend maintained post-HX (Figure 7K-M). Since *Sparc* encodes a secreted protein that negatively regulates synaptogenesis^62^, this emphasizes the specialized function of cortical astrocytes in synapse formation and neural homeostasis.

OL-lineage cells in different regions showed different levels of gene expression for OL differentiation and maturation related genes. OPCs in the SVZ and corpus callosum had higher levels of oligodendrogenesis genes including *Olig2*, *Gpr17*, *Itpr2*, and *Fyn* compared to cortical OPCs (Figure 7A-B, 7D-E). These genes were upregulated in SVZ OPCs following HX, while only some (i.e., *Gpr17*) were upregulated in corpus callosum OPCs (Figure 5H, Table S2). To characterize the overall differentiation and maturation state of OL-lineage cells, we used several established genes^63–65^ (see *Methods*) to calculate scores for OL differentiation priming (“OL genesis”), and OL maturation. We found that OPCs in the SVZ and corpus callosum had a higher OL genesis score than both superficial and deep layer cortical OPCs. Similarly, newly formed OLs had a higher OL genesis score in the corpus callosum than in the superficial and deep cortical layers. Following HX, there was a notable increase in the OL genesis scores among OPCs in the SVZ and corpus callosum, as well as in newly formed OLs in the corpus callosum (Figure 7F-G). The OL maturation score was higher in immature and mature OLs from the corpus callosum than in those in the superficial and deep cortical layers. This score increased significantly in upper cortical OLs following HX (Figure 7H-J). Overall, these results suggest an adaptive response to injury involving an increase in the generation and maturation of OL lineage cells, and a regionalization in the maturation dynamics between corpus callosum and cortical OLs.

## DISCUSSION

Advances in the care of very low birthweight preterm infants have markedly improved survival. However, despite these advances, survivors of preterm birth continue to have high propensity for developmental delay and disability. Chronic HX exposure resulting from an underdeveloped respiratory system leads to cerebral WMI, a major cause of neurodevelopmental disorders in preterm infants, including cerebral palsy. However, the underlying cellular mechanisms that govern the brain’s response to HX and recovery from WMI remain poorly understood. Here, we employed high-resolution spatial transcriptomics as a discovery platform to map cell type- and region-specific transcriptional signatures during recovery from perinatal HX. At baseline (NX condition), we identified several brain region-specific cell states, including spatially restricted populations of glia and vascular cells that play a key role in modulating the signaling environment and may regulate developmental processes such as neurogenesis and gliogenesis, cell migration, synapse formation, and inflammation. We found that during the reparative phase following neonatal HX exposure, there were changes in regional cell type composition, neurogenic and gliogenic transcriptional programs, and local cellular architecture, which are likely the result of extensive changes in regional signaling networks. Thus, our results support a model of niche cell states and signaling networks that are region- and stimulus-dependent.

First, we demonstrate that at baseline, OPCs in the neurogenic niche and white matter have a transcriptomic profile that is more proliferative and more primed for differentiation than in the cortex. In response to HX, OPCs within the neurogenic niche further upregulate proliferation-associated genes and genes encoding ligands that modulate OL formation and myelination. We found that OPCs upregulated BMP ligand genes such as *Bmp4* which, in the context of demyelinating injury, has been shown to be produced by injury-activated OPCs and inhibit the differentiation of OPCs to OLs.^66^ We also observed upregulation of *Cx3cl1*, which encodes fractalkine, from several glial cell types including OL-lineage cells to reactive microglia. Fractalkine has an established pro-regenerative role in remyelination, particularly via OPCs and microglia.^67,68^ Interestingly, previous work has shown that environmental enrichment both enhances recovery from neonatal diffuse WMI and increases fractalkine levels. Importantly, TGFβ is known to regulate fractalkine expression, and we identified increased TGFβ1 signaling among glial and vascular cell types, which likely acts as an upstream regulator of tissue remodeling after HX. While OPCs in the white matter became more abundant, newly formed OLs became less abundant, suggesting differentiation arrest, specifically in the transition from OPCs to newly formed OLs.

Further, we found that developing and mature OLs from the cortex and corpus callosum exhibited different cell states and responded differently to HX. At baseline, white matter OLs overall were more mature than cortical OLs (exhibiting higher OL maturation scores), and these levels did not change significantly following HX. Analysis of inter-lineage cell-cell communication revealed alterations in GAS6-TAM, EDN1-EDNRB, and fractalkine-CX3CR1 networks in the cortex that may play a direct role in promoting OPC differentiation and altering OL myelination capacity in the context of neonatal HX. In addition, we identify significant changes in the spatial organization of homeostatic astrocytes. Previous studies have shown that astrocyte-OL interactions are crucial in facilitating OPC differentiation into mature myelin-forming OL via a process orchestrated through the Nrf2 pathway.^69^

Importantly, we identified profound and widespread changes in extracellular matrix components, including CSPGs, after HX. CSPGs contribute to the formation of perineuronal nets (PNNs) during postnatal development and play a role in the closure of the critical window of plasticity.^70,71^ We observed that in most cell types in the cortex and corpus callosum, perinatal HX was associated with increased expression of genes important for cell migration and projection guidance including ephrins and CSPGs. These signals may play an important role in coordinating the migration of injury-responsive cells such as immune cells and astrocytes, as well as migrating upper-layer excitatory neurons. Interestingly, we did find that the abundance and local organization of upper-layer excitatory neurons was disrupted by HX. However, it is unclear whether the upregulation of guidance cues such as CSPGs would help or hinder appropriate cell migration. Depletion of CSPGs in the adult brain by enzymatic degradation or genetic inhibition has been shown to promote plasticity.^72–74^ CSPGs can also act as potent inhibitors of myelination, axon guidance and regeneration, and other injury repair processes. Further work is needed to determine the clinical relevance of CSPGs in myelin recovery after perinatal WMI.

While most studies on neonatal WMI have focused on OL-lineage cells, our study underscores the complex and region-specific multicellular signaling that may support gliogenesis but also ultimately lead to impaired functional connectivity. Specifically, we found that endothelial cells, fibroblasts, and pericytes were major sources of signaling molecules, and that their phenotype changed dramatically following HX in a region-specific manner. Our findings imply that crosstalk signaling from vasculature-associated cells may be crucial for activating angiogenesis and forming adhesive interactions that maintain blood-brain barrier integrity and modulate permeability after HX exposure. Additionally, vascular cells may represent an important source of inflammatory ligands that promote microglial reactivity after perinatal HX. We also found that corpus callosum microglia may be less reactive than cortical microglia under NX and HX conditions, which contrasts with previous research in adult rodents and humans, that suggest that white matter microglia are more reactive to injury than gray matter microglia.^75,76^ As these earlier studies primarily involved adult and aged rodents, our findings reflect a different, developmental microglia state. Specifically, the heightened cortical microglial reactivity observed in our study may relate to their role in synapse refinement during brain development following HX exposure, although this requires further investigation.

An important strength of the present study is the use of MERFISH, which enables single cell-resolution spatial transcriptomics. Previous work on neonatal HX has focused largely on the corpus callosum, but paracrine and non-cell autonomous mechanisms of recovery after neonatal brain injury are poorly understood. The present analysis spans most major anatomical brain regions and highlights regional variability in the vulnerability and response to injury. Nonetheless, there are several limitations to the present study. Firstly, the mouse model of chronic neonatal HX does not introduce inflammatory stimuli, which are often an important component of the encephalopathy of prematurity. Additionally, the sub-lethal chronic HX model employed does not fully replicate the intermittent hypoxic events experienced by many premature infants. Notably, the pattern of WMI in preterm infants has become less severe in the last two decades, and thus, cerebral dysmaturation may also be contributing to neurodevelopmental delays and impairments.^77^ From a methodologic perspective, MERFISH requires segmentation of cell boundaries to assign transcripts to individual cells. We employed a machine learning algorithm and extensive manual curation, including stringent doublet elimination, to mitigate limitations of cell boundary identification. However, precise cell boundary identification remains an ongoing challenge for *in situ* spatial transcriptomics, and mis-segmentation may introduce false cell type assignment of gene expression. While our analysis was limited to a single time point, the objective of the study was to identify pathways critical for recovery and repair after perinatal HX. Finally, commercial MERFISH methods are presently limited to 500 genes, so additional changes at the level of the whole transcriptome were not captured.

Despite these limitations, this work presents an extensive spatially informed landscape of gene expression changes in an established mouse model of perinatal brain injury. We developed and employed a new approach to map physical cellular interactions using imaging-based spatial transcriptomics, which can be extended to other developmental and disease contexts. Our whole-brain analysis is a foundational resource for developmental WMI and for regenerative cellular communication.

## Resource Availability

### Lead Contact

Further information and requests for resources and reagents should be directed to and will be fulfilled by the lead contact, Brian T. Kalish

### Materials availability

This study did not generate new unique reagents.

### Data and code availability

MERFISH data have been deposited in GEO under accession code **GSE255892**. All data generated or analyzed during this study are included in the manuscript and supporting files.

The code for differential proximity analysis is available on Github at: https://github.com/keon-arbabi/spatial_hypoxia/blob/main/keon_spatial_analyses.R

### AUTHOR CONTRIBUTIONS

Overall conceptualization, B.T.K. and S.J.T; tissue acquisition and processing, B.T.K. and N.T.; MERFISH sample preparation and imaging, N.T.; computational data analysis, N.T., M.Y.F, K.A., R.X., B.K., M.W., S.J.T., and B.R.; manuscript preparation and writing, B.T.K., N.T., M.Y.F., K.A., S.J.T, A.L., and B.R.; figure design and preparation, B.R.; funding acquisition, B.T.K.; supervision and project administration, B.T.K.

## METHODS

### Mouse Model of Chronic Sublethal Hypoxia

All animal experiments were performed in accordance with the Canadian Council of Animal Care policies. CD-1 mice (Strain: 022) were purchased from Charles River Laboratories. The animals were kept in 12/12 hour light/dark cycles with free access to food and chow, and bred at the SickKids Research Institute in Toronto (ON, Canada). Offspring CD-1 mice were exposed to an oxygen concentration of either 10% (HX) or 21% (NX) in a HX chamber from postnatal day 3 (P3) to P11, lasting eight consecutive days.^7,9,78^ Mouse dams were rotated daily. At P11, mice were removed from the chamber and transferred to a room with normoxic conditions (21% oxygen) until euthanasia by CO_2_ at P21. Fresh brain tissue was harvested and immediately placed in optimal cutting temperature (OCT) solution and frozen at -80°C until MERFISH processing. Male offspring were used for MERFISH experiments.

### MERFISH Gene Panel Design

We created a panel of 497 genes (Table S1) to study the cellular architecture and signaling landscape of the murine forebrain using MERFISH. To guide the identification of diverse cell types in the brain, the panel included 181 genes that are known cell-type specific markers or have been previously shown to have enriched expression in a specific population of cells. This list targeted known cell types within major forebrain structures including the cortex, white matter tracts, striatum, and SVZ. The remaining 316 genes were components of biological pathways shown previously or hypothesized to play a role in the pathophysiology of chronic neonatal HX. This included components of signaling pathways known to regulate oligodendrogenesis, myelination, or that have previously been implicated in the pathophysiology of WMI such as WNT signaling^79,80^, BMP signaling^81,82^, SHH signaling^82,83^, FGF signaling^84,85^, EGF signaling^86^, Ephrin signaling^51,87^, cytokine signaling^88^, senescence^89,90^, and oncogenes.^91^ Each gene was assigned a 20-bit binary barcode which was used to decode sequential imaging data to assign detected transcripts to the appropriate gene.

### MERFISH Tissue Processing and Sample Preparation

All MERFISH sample preparation was performed under RNase-free conditions. Frozen embedded brains were cryo-sectioned at 10 μm thickness at -21°C, and mounted onto room temperature (RT) MERSCOPE beaded coverslips (Vizgen, Cat: 10500001). After adhering, sections were allowed to refreeze for 5-15 min, then fixed in 4% paraformaldehyde (PFA) diluted in 1X PBS for 15 min. Sections were washed three times with 1X PBS for 5 min each, then stored in 70% ethanol (EtOH) overnight at 4°C to permeabilize the tissue. Sections were stored in 70% EtOH for a maximum of 3 weeks.

Sample preparation was performed using the sample preparation kit (Vizgen, Cat: 10400012) and Vizgen manufacturer instructions for unfixed tissue. First, sections were washed with 1X PBS followed by Sample Prep Wash Buffer (Vizgen, PN 20300001). Sections were incubated in Formamide Wash Buffer (Vizgen, PN 20300002) for 30 min at 37°C, then incubated in the gene panel mix for 42-46 hour (hr) at 37°C. Sections were then incubated two times in Formamide Wash Buffer for 30 min each at 47°C, and washed with sample prep wash buffer for at least 2 min. Sections were coated in gel embedding solution (0.05% w/v ammonium persulfate, 0.05% v/v N,N,N’,N’-tetramethylethylenediamine in Gel Embedding Premix (Vizgen, PN 20300004)) and incubated at RT for 1.5 hr, cleared in 1:100 proteinase K in Clearing Premix (Vizgen, PN 20300003) at 37°C overnight or for a maximum of 7 days to clear lipids and proteins that may contribute to auto-fluorescence background noise. Prior to imaging, sections were washed two times with Sample Prep Wash Buffer, incubated in DAPI and PolyT Staining Reagent (Vizgen, PN 20300021) for 15 min on a rocking platform, incubated in Formamide Wash Buffer for 10 min, and washed again with Sample Prep Wash Buffer.

### MERFISH Imaging

Imaging was performed on the MERSCOPE platform (Vizgen, Cat: 10000001) according to manufacturer instructions. Briefly, samples were loaded into the flow chamber of the instrument and the desired region for imaging was selected using a low-resolution mosaic of DAPI and PolyT stains. Samples were imaged at high-resolution with a 7-plane *z*-stack and 1.5 μm spacing between adjacent *z*-planes to capture the entire 10 μm thickness of the tissue sections. Samples were then automatically imaged according to MERSCOPE imaging presets.

### MERFISH Bioinformatics Workflow

After imaging, transcript barcodes were decoded and assigned to the appropriate gene. MERFISH images were segmented using Vizgen’s post-processing tool (VPT) and Cellpose 2.0^22^, a machine learning algorithm (RRID:<u>SCR_021716</u>). DAPI and PolyT signals were used to delineate cell boundaries for each field-of-view. Individual RNA molecules were assigned to a cell based on whether they were positioned within the marked boundary. Anatomical segmentation was performed based on tissue morphology and region-specific gene expression patterns. This revealed 8 gross anatomical regions: cortex (marked by high expression of *Slc7a6* and *Slc17a7*); corpus callosum, anterior commissure, septal white matter tracts (all marked by high expression of *Cnp* and *Plp1*), SVZ (marked by high expression of *Foxj1*, *Sox2*, and *Ascl1*), caudoputamen (marked by high expression of *Foxo1*, *Crym*, and *Gad1/2)*, lateral septal nucleus (marked by expression of *Sp8* and *Sp9*), anterior commissure (marked by high expression of *Cnp*, and *Plp1*), bed nucleus of the stria terminalis (marked by *Cartpt*), and the regions ventral to the caudoputamen and anterior commissure, which were collectively labeled lower gray matter (high expression of *Gad1*, and *Dlx1/2*). Meninges (high expression of *Col1a1* and *Gfap*) were grouped with the cortex. The cortex was further segmented into upper, superficial cortex layers (marked by high expression of *Cux2*, *Cartpt*, *Kitl*, and *Rgs8*), and deeper cortex layers (marked by high expression of *Bcl11b*, *Fezf2*, *Otx1*, and *Tle4*).

To obtain a cell-by-gene matrix of each biological sample tissue replicate, we adapted an existing bioinformatic pipeline^19^ as follows. (1) To remove segmentation artifacts of extremely small or large cells, we removed cells with a volume less than 50µm^3^ or larger than three times the median volume of all cells. (2) Cells with zero RNA molecules were removed. (3) In a 10µm^2^ thick tissue slice, the spatial orientation of cells within the section resulted in partial imaging of their soma. To account for potential RNA discrepancies, gene expression for each cell was normalized by their volume and multiplied by 1000. (4) Cells with total RNA counts falling below 2% quantile or exceeding the 98% quantile were excluded. (5) Potential doublets were removed using Scrublet (RRID:<u>SCR_018098</u>), a program that generates artificial doublets by comparing gene expression profiles of randomly selecting cells with segmented cells in the dataset and using a k-nearest neighbor (kNN) to output a predicted doublet score. Cells with a Scrublet-based doublet score of greater than 0.20 were excluded for further analysis, accounting for 2.5-3.5% of cells across samples.

Data was then processed using the Seurat V5 package in R (RRID:<u>SCR_016341</u>). First, each dataset was normalized using Seurat’s SCTransform function and cells from each gross anatomical region were grouped into separate Seurat objects. We focused subsequent analyses on cells contained in 3 brain regions: cortex, corpus callosum, and SVZ. We integrated and scaled gene expression from cells from each of six samples (*n* = 3 HX and *n* = 3 NX). Only genes with sufficient expression across cells in that region (ranging from 475-495 genes) were integrated and used to compute principal components and uniform manifold approximation and projection (UMAP). Cell clustering was performed using shared nearest neighbors-cliq (SNN-cliq). Cell typing and subtyping was performed using canonical markers. To improve our cell type identification, we removed (1) clusters that did not express cell type-specific marker genes, and thus could not be assigned to a cell type, and (2) clusters of cells that expressed two or more mutually exclusive cell-type-specific marker genes. The former excluded potential non-cell artifacts or cells which could not be identified with the implemented gene panel, and the latter excluded potential doublets.

### Cell Proportion Analysis

To determine differences in Cell Type abundance in response to HX, we calculated cell type proportions for each Cell SubType class within each region. For each biological replicate and each region, the total number of cells in each Cell SubType was divided by either the total number of cells in the region or the total number of cells in the region corresponding to a broad cell class (e.g., neurons, glia, OL-lineage). Statistical differences in cell proportions were calculated using an unpaired, two-tailed, Student’s t-test comparing HX and NX (*n* = 3 each) for each Cell SubType of each region.

### Cell-Level Proximity Analysis

To understand how the composition of cellular neighborhoods are affected by perinatal HX, we developed a novel method to quantify condition-related differences in local cell type proximity. For each cell *i*, a k-dimensional tree was used to count the number of neighboring type B cells, *N_B_*, and the total number of neighboring cells (irrespective of type), *N_total_*, within a specified radius *r*. For each sample and brain region, we defined a minimum distance unit, *d*, as the median of the smallest distances between all cell pairs for each sample. A restricted radius (*r* = 5 * *d*) was selected to focus on the immediate proximity interactions between Cell Types. The proportion of type B cells within this radius is given by:

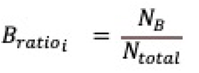

The *B_ratio_* provides a normalized measure of the local proportion or density of cell type B in the proximity to a reference cell *i*. We note that *B_ratio_* is asymmetric (e.g., the proportion of astrocytes in the proximity of neurons is different from the proportion of neurons in the proximity of astrocytes), which allows for discriminating when the neighborhood of one cell type is enriched or depleted for another cell type, but not vice versa. We further note that this analysis of local cell proximity or density is conceptually similar to the analysis of global cell proportion differences defined above.

To identify differences in local cell type proximity under NX and HX conditions, we used a Generalized Linear Mixed-Effects Model (GLMM) implemented via the glmmTMB R package and goodness-of-fit calculations evaluated by the DHARMa package (RRID:<u>SCR_022136</u>). This statistical model addresses the non-independence of within-tissue sample observations and is applied to the log-transformed *B_ratio_* , with a small offset to accommodate zeroes. This approach quantifies the impact of condition on cell type distributions, while accounting for inter-sample variability:

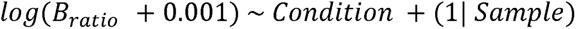

The effect size and FDR for the condition coefficient were assessed. Significant coefficients (FDR < 0.05) indicated a change in cell proximity under HX conditions relative to NX, suggesting a disruption in cellular neighborhoods. A positive effect size indicated that cell type B is more densely clustered near cell type A (center cell) in HX compared to NX, suggesting increased proximity. In contrast, a negative effect size indicated that cell type B is closer to cell type A in NX compared to HX, indicating a relative decrease in proximity under hypoxic conditions.

### Differential Gene Expression Analysis

To identify the differentially expressed genes between conditions and regions, we employed MAST^92^ using Seurat’s FindMarkers function on the raw counts matrix. Only genes expressed in a minimum of 5% of cells in either comparison group were tested. P-values were adjusted with the Benjamini-Hochberg (BH) correction method to obtain FDR values. Specific log_2_(FC) values are reported in Table S2. Genes with a |log_2_(FC)| > 0.25 and FDR < 0.05 were considered significantly differentially expressed.

### Cell State Scoring

To calculate cell state scores, we selected genes whose expression has been previously shown to be enriched in cells primed for OL differentiation (OL genesis score) or enriched in mature OLs (OL maturation score^63–65^; enriched in reactive astrocytes over homeostatic astrocytes (reactive astrocyte score^59–61^); and enriched in reactive microglia compared to homeostatic microglia (reactive microglia score).^57^ Cell state scores were calculated for each cell from the normalized, integrated datasets by taking the average *z*-score for the genes in the cell state gene list in each cell. Then, cell state scores for specific cell groups (i.e., specific Cell Types or Cell SubTypes for each replicate for each anatomical region) were calculated by taking the average cell state score across all cells within that group of cells. To compare changes from NX to HX, statistical analysis was performed using the two-tailed, unpaired, Student’s t-test comparing the cell state scores from the 3 NX replicates to 3 HX replicates within each cell type and anatomical region analyzed. Genes used for the OL genesis score were *Ptprz1*, *Qk*, *Itpr2*, *Gpr17*, *Fyn*, and *Tcf7l2*. Genes used for the OL maturation score were *Mog*, *Mag*, *Cnp*, *Plp1*, *Myrf*, *Egr2*, *Fos*, *Fosb*, *Klk6*, *Ptgds*, *Car2*, *Grm3*, *Npsr1*, *Jph4*, *Aspa*, *Msmo1*, *Sqle*, *Hmgcs1*, *Idi1*, *Opalin*, and *Trf*. Genes used for the reactive astrocyte score were *Serpina3n*, *C3*, *Emp1*, *Gfap*, *Ggta1*, *H2-T23*, *Cd109*, *Hspb1*, and *lcn2*. Genes used for the reactive microglia score were *Lpl*, *Cst7*, *Ptprc*, *Trem2*, *Lgals3*, *Axl*, *Lyz2*, *Clec7a*, *Spp1*, *Cst7*, and *Apoe*.

### Ligand-Receptor (LR) Analysis

We performed ligand-receptor (LR) interaction analysis on the raw gene expression datasets using CellChat v2 (RRID:SCR_021946).^93^ For each experimental condition, spatial coordinates were aggregated, and an average spot size was computed to normalize the interaction distances.

Coordinates from individual replicates were appropriately translated to prevent overlap and maintain the spatial integrity of the samples. The preprocessed data was then converted to a CellChat object, which encapsulated the gene expression matrix, cellular metadata, and spatial coordinates. In our analysis, we grouped cells based on both Cell Type and Cell SubType classes and incorporated spatial data to contextualize cell interactions within the tissue architecture. We then aligned our gene expression data with established molecular communication networks to predict potential LR and in turn cell-cell interactions. The likelihood and extent of intercellular signaling interactions was estimated by assessing the expression of LR pairs. Truncated mean calculation for average gene expression per cell group and bootstrapping for statistical inference were performed to ensure estimation robustness. The communication probabilities were calculated, accounting for the effect of spatial distance on signaling potential by adjusting for the scaled interaction range and considering contact-based nearest neighbor interactions. Subsequently, pathways were aggregated, and centrality analysis was performed to identify key nodes within the communication network. To explore the differential intercellular communication networks between HX and NX, data from both conditions were integrated. Differential expression analysis was then carried out to identify genes that displayed significant changes in expression levels, with a threshold of p-value < 0.01. Changes in ligand and receptor expression level, with a |log_2_(FC)| > 0.25 were considered significant.

## Supplemental Information

**Figure S1:**
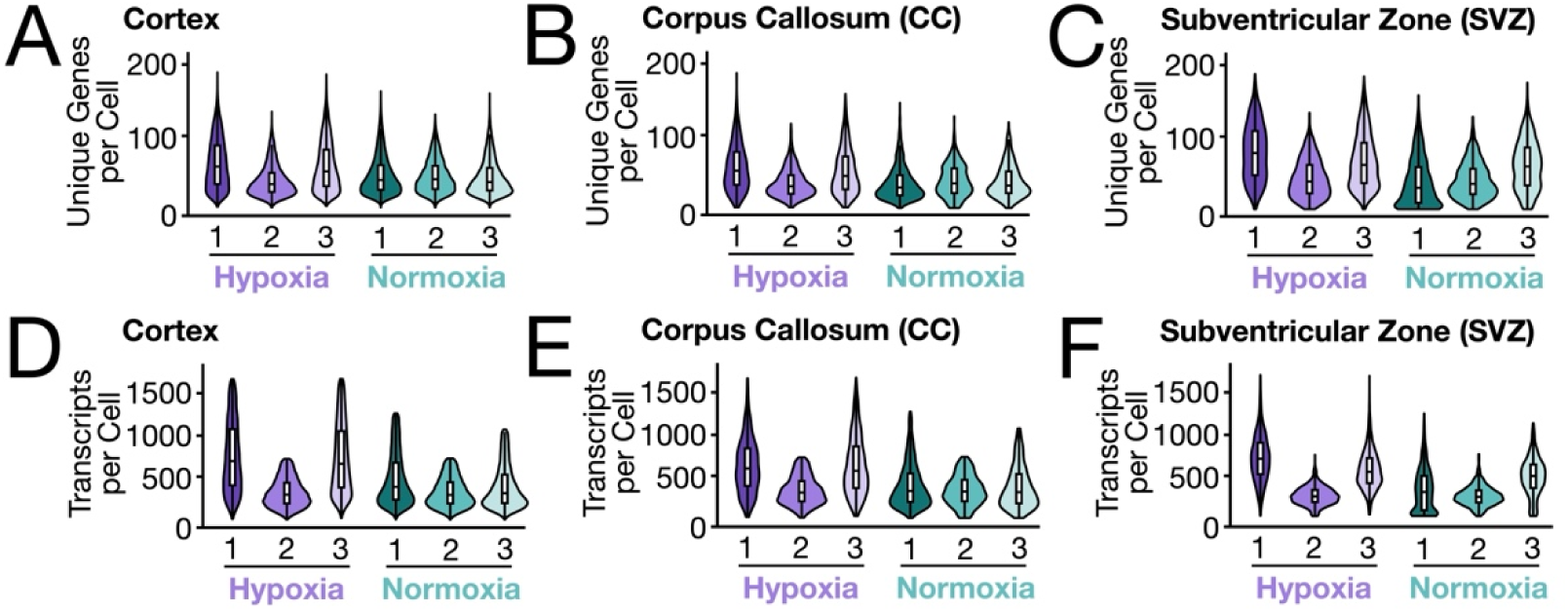
Quality control metrics of the MERFISH data. Violin plots showing select quality control parameters used to profile the data, including the number of unique genes per cell in the **(D)** cortex, **(E)** corpus callosum, and **(F)** SVZ, and the number of transcripts per cell in the **(G)** cortex, **(H)** corpus callosum, and **(I)** SVZ.

**Figure S2:**
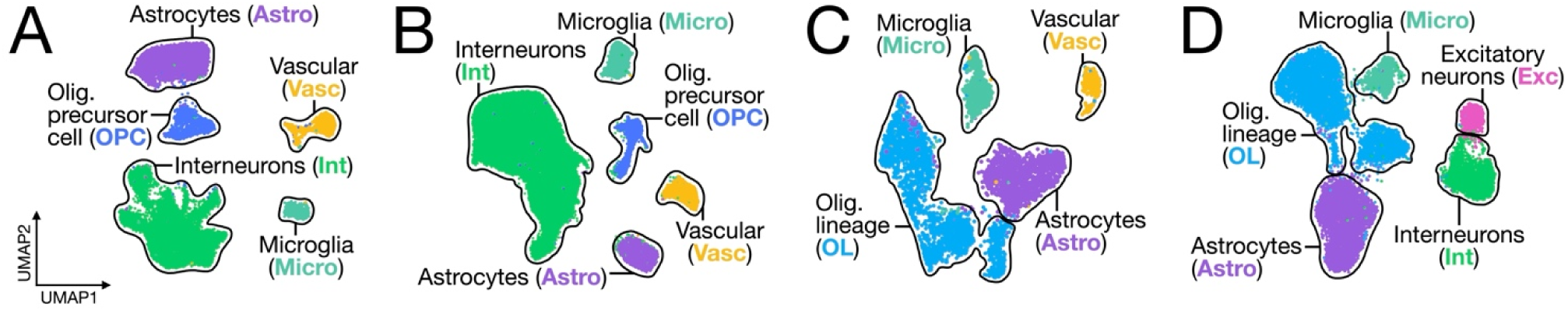
Cell Type identification across the P21 brain. UMAP visualization of cell types identified in the **(A)** septum, **(B)** caudate putamen (CP), **(C)** anterior commissure (ACo), and **(D)** septal white matter tracts (SWM), where each dot represents a single cell. UMAP plots are generated from combined replicates across NX and HX conditions. Each cluster is colored by Cell Type.

**Figure S3:**
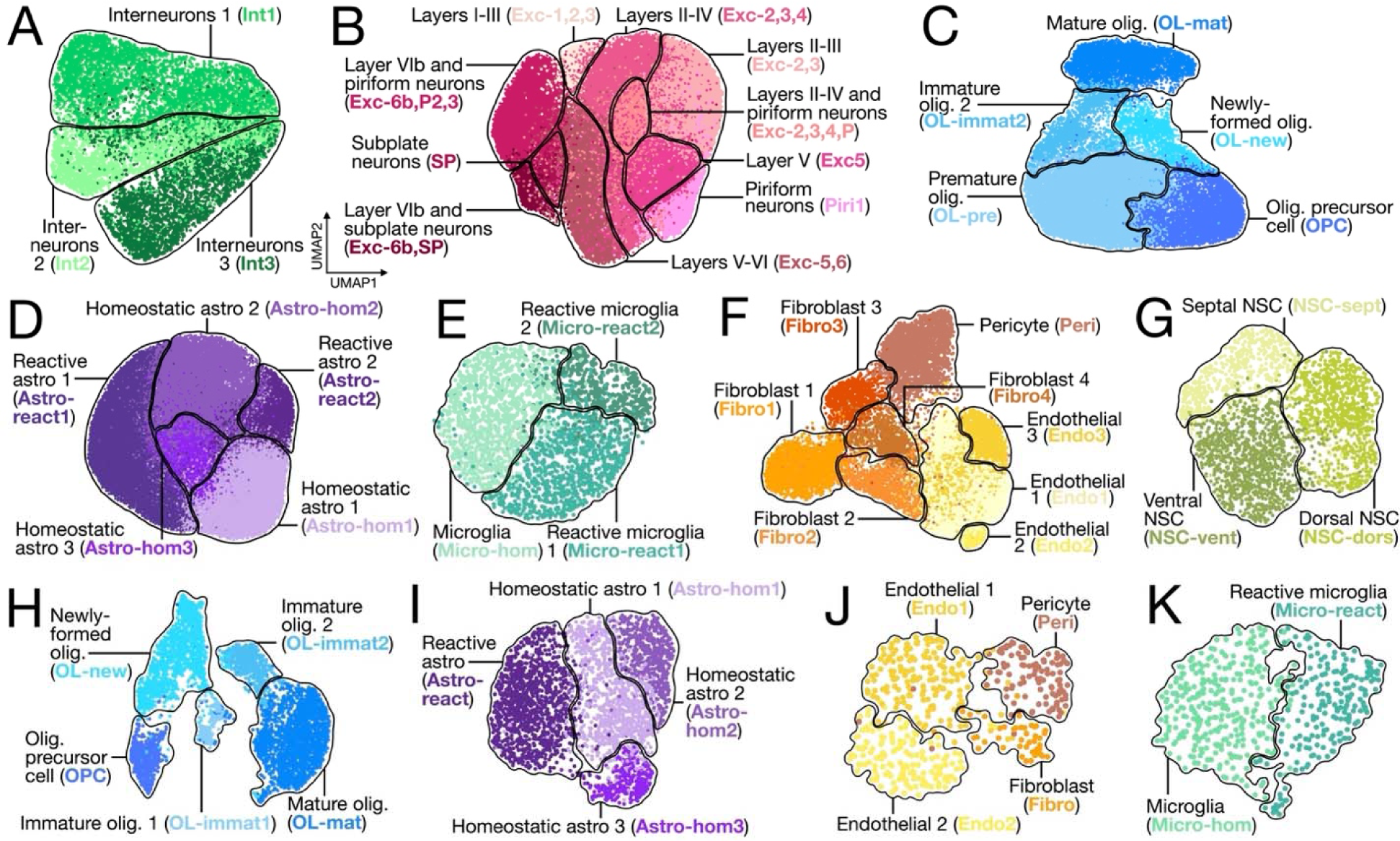
Cell Subtype identification across the cortex, SVZ and corpus callosum of the P21 brain. UMAP visualization of cortical **(A)** interneurons and **(B)** excitatory neurons, **(C)** OL-lineage cells, **(D)** astrocytes, **(E)** microglia, and **(F)** vascular cells; **(G)** SVZ neural stem cells (NSC); and corpus callosum **(H)** OL-lineage cells, **(I)** astrocytes, **(J)** vascular cells, and **(K)** microglia, where each dot represents a single cell. UMAP plots are generated from combined replicates across NX and HX conditions. Each cluster is colored by Cell Subtype as previously defined.

**Figure S4:**
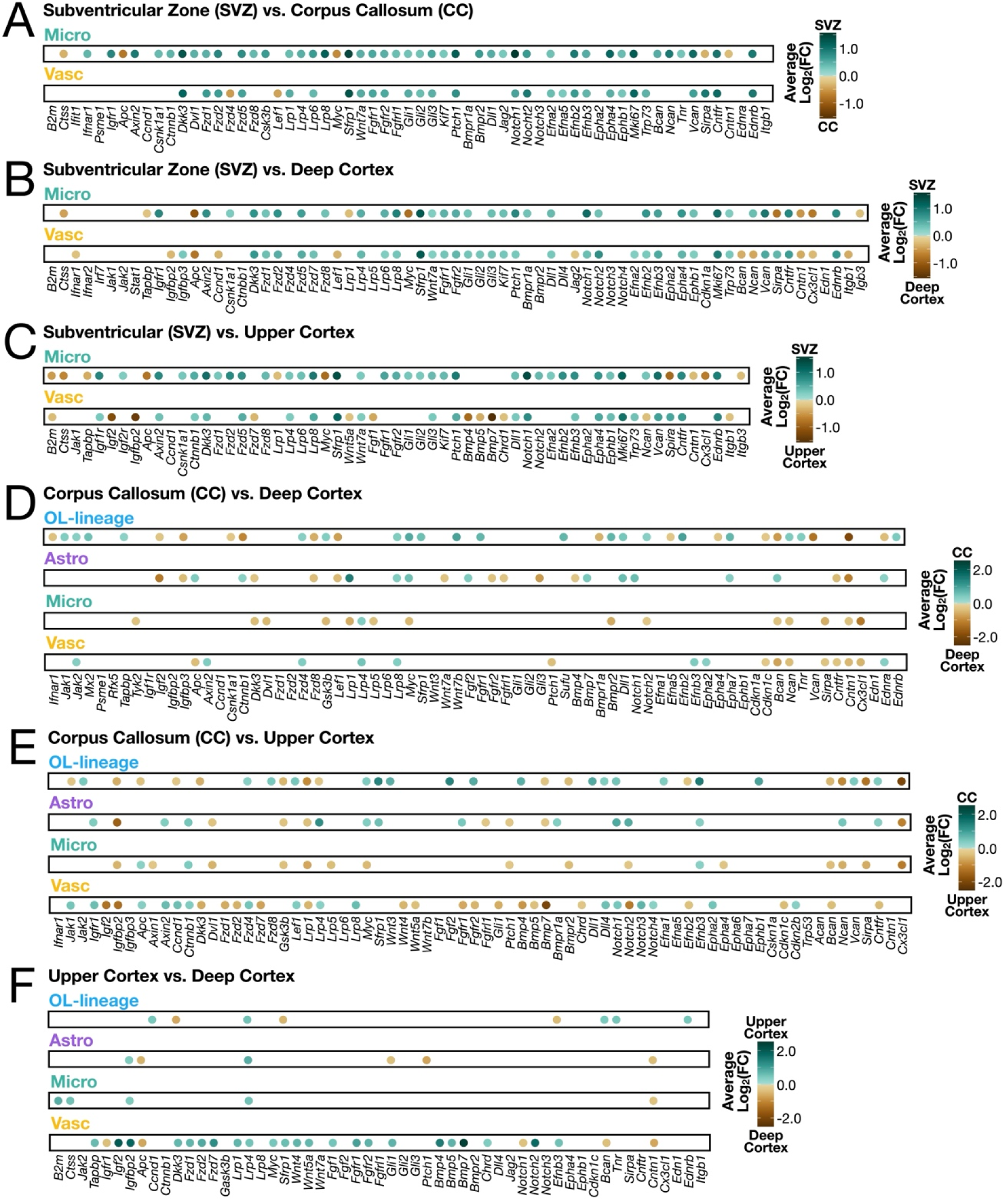
Comparison of signaling-related gene expression between anatomical regions in mice exposed to normoxic conditions. Dotplots displaying differential gene expression results comparing Cell Types between SVZ, corpus callosum, deep cortex, and upper cortex within NX mice. Only significant genes (FDR < 0.05) are shown. The color of the dot represents the average log_2_(FC) as per each associated legend. Comparisons shown are **(A)** SVZ versus corpus callosum, **(B)** SVZ and deep cortex, **(C)** SVZ versus upper cortex, **(D)** corpus callosum versus deep cortex, **(E)** corpus callosum versus upper cortex, and **(F)** deep cortex versus upper cortex. Results are also shown in Table S2.

## REFERENCES

1. Blencowe, H., Cousens, S., Chou, D., Oestergaard, M., Say, L., Moller, A.-B., Kinney, M., Lawn, J., and Born Too Soon Preterm Birth Action Group (2013). Born too soon: the global epidemiology of 15 million preterm births. Reprod. Health 10 *Suppl 1*, S2. 10.1186/1742-4755-10-S1-S2.

2. Blencowe, H., Lee, A.C.C., Cousens, S., Bahalim, A., Narwal, R., Zhong, N., Chou, D., Say, L., Modi, N., Katz, J., et al. (2013). Preterm birth-associated neurodevelopmental impairment estimates at regional and global levels for 2010. Pediatr. Res. 74 *Suppl 1*, 17–34. 10.1038/pr.2013.204.

3. Crump, C., Sundquist, J., and Sundquist, K. (2021). Preterm or Early Term Birth and Risk of Autism. Pediatrics 148, e2020032300. 10.1542/peds.2020-032300.

4. Rommel, A.-S., James, S.-N., McLoughlin, G., Brandeis, D., Banaschewski, T., Asherson, P., and Kuntsi, J. (2017). Association of Preterm Birth With Attention-Deficit/Hyperactivity Disorder-Like and Wider-Ranging Neurophysiological Impairments of Attention and Inhibition. J. Am. Acad. Child Adolesc. Psychiatry 56, 40–50. 10.1016/j.jaac.2016.10.006.

5. Hirvonen, M., Ojala, R., Korhonen, P., Haataja, P., Eriksson, K., Gissler, M., Luukkaala, T., and Tammela, O. (2017). The incidence and risk factors of epilepsy in children born preterm: A nationwide register study. Epilepsy Res. 138, 32–38. 10.1016/j.eplepsyres.2017.10.005.

6. Salmaso, N., Jablonska, B., Scafidi, J., Vaccarino, F.M., and Gallo, V. (2014). Neurobiology of premature brain injury. Nat. Neurosci. 17, 341–346. 10.1038/nn.3604.

7. Jablonska, B., Adams, K.L., Kratimenos, P., Li, Z., Strickland, E., Haydar, T.F., Kusch, K., Nave, K.-A., and Gallo, V. (2022). Sirt2 promotes white matter oligodendrogenesis during development and in models of neonatal hypoxia. Nat. Commun. 13, 4771. 10.1038/s41467-022-32462-2.

8. Forbes, T.A., Goldstein, E.Z., Dupree, J.L., Jablonska, B., Scafidi, J., Adams, K.L., Imamura, Y., Hashimoto-Torii, K., and Gallo, V. (2020). Environmental enrichment ameliorates perinatal brain injury and promotes functional white matter recovery. Nat. Commun. 11, 964. 10.1038/s41467-020-14762-7.

9. Fagel, D.M., Ganat, Y., Silbereis, J., Ebbitt, T., Stewart, W., Zhang, H., Ment, L.R., and Vaccarino, F.M. (2006). Cortical neurogenesis enhanced by chronic perinatal hypoxia. Exp. Neurol. 199, 77–91. 10.1016/j.expneurol.2005.04.006.

10. Herrero-Navarro, Á., Puche-Aroca, L., Moreno-Juan, V., Sempere-Ferràndez, A., Espinosa, A., Susín, R., Torres-Masjoan, L., Leyva-Díaz, E., Karow, M., Figueres-Oñate, M., et al. (2021). Astrocytes and neurons share region-specific transcriptional signatures that confer regional identity to neuronal reprogramming. Sci. Adv. 7, eabe8978. 10.1126/sciadv.abe8978.

11. Joglekar, A., Prjibelski, A., Mahfouz, A., Collier, P., Lin, S., Schlusche, A.K., Marrocco, J., Williams, S.R., Haase, B., Hayes, A., et al. (2021). A spatially resolved brain region- and cell type-specific isoform atlas of the postnatal mouse brain. Nat. Commun. 12, 463. 10.1038/s41467-020-20343-5.

12. Tan, Y.-L., Yuan, Y., and Tian, L. (2020). Microglial regional heterogeneity and its role in the brain. Mol. Psychiatry 25, 351–367. 10.1038/s41380-019-0609-8.

13. Stogsdill, J.A., Kim, K., Binan, L., Farhi, S.L., Levin, J.Z., and Arlotta, P. (2022). Pyramidal neuron subtype diversity governs microglia states in the neocortex. Nature 608, 750–756. 10.1038/s41586-022-05056-7.

14. Spitzer, S.O., Sitnikov, S., Kamen, Y., Evans, K.A., Kronenberg-Versteeg, D., Dietmann, S., de Faria, O., Agathou, S., and Káradóttir, R.T. (2019). Oligodendrocyte Progenitor Cells Become Regionally Diverse and Heterogeneous with Age. Neuron 101, 459–471.e5. 10.1016/j.neuron.2018.12.020.

15. Hilscher, M.M., Langseth, C.M., Kukanja, P., Yokota, C., Nilsson, M., and Castelo-Branco, G. (2022). Spatial and temporal heterogeneity in the lineage progression of fine oligodendrocyte subtypes. BMC Biol. 20, 122. 10.1186/s12915-022-01325-z.

16. Brandi, E., Torres-Garcia, L., Svanbergsson, A., Haikal, C., Liu, D., Li, W., and Li, J.-Y. (2022). Brain region-specific microglial and astrocytic activation in response to systemic lipopolysaccharides exposure. Front. Aging Neurosci. 14, 910988. 10.3389/fnagi.2022.910988.

17. Hammond, T.R., Dufort, C., Dissing-Olesen, L., Giera, S., Young, A., Wysoker, A., Walker, A.J., Gergits, F., Segel, M., Nemesh, J., et al. (2019). Single-Cell RNA Sequencing of Microglia throughout the Mouse Lifespan and in the Injured Brain Reveals Complex Cell-State Changes. Immunity 50, 253–271.e6. 10.1016/j.immuni.2018.11.004.

18. Zhang, M., Eichhorn, S.W., Zingg, B., Yao, Z., Cotter, K., Zeng, H., Dong, H., and Zhuang, X. (2021). Spatially resolved cell atlas of the mouse primary motor cortex by MERFISH. Nature 598, 137–143. 10.1038/s41586-021-03705-x.

19. Allen, W.E., Blosser, T.R., Sullivan, Z.A., Dulac, C., and Zhuang, X. (2023). Molecular and spatial signatures of mouse brain aging at single-cell resolution. Cell 186, 194–208.e18. 10.1016/j.cell.2022.12.010.

20. Zhang, M., Pan, X., Jung, W., Halpern, A.R., Eichhorn, S.W., Lei, Z., Cohen, L., Smith, K.A., Tasic, B., Yao, Z., et al. (2023). Molecularly defined and spatially resolved cell atlas of the whole mouse brain. Nature 624, 343–354. 10.1038/s41586-023-06808-9.

21. Semple, B.D., Blomgren, K., Gimlin, K., Ferriero, D.M., and Noble-Haeusslein, L.J. (2013). Brain development in rodents and humans: Identifying benchmarks of maturation and vulnerability to injury across species. Prog. Neurobiol. 106–107, 1–16. 10.1016/j.pneurobio.2013.04.001.

22. Pachitariu, M., and Stringer, C. (2022). Cellpose 2.0: how to train your own model. Nat. Methods 19, 1634–1641. 10.1038/s41592-022-01663-4.

23. Hutter, G., Theruvath, J., Graef, C.M., Zhang, M., Schoen, M.K., Manz, E.M., Bennett, M.L., Olson, A., Azad, T.D., Sinha, R., et al. (2019). Microglia are effector cells of CD47-SIRPα antiphagocytic axis disruption against glioblastoma. Proc. Natl. Acad. Sci. U. S. A. 116, 997–1006. 10.1073/pnas.1721434116.

24. Ding, X., Wang, J., Huang, M., Chen, Z., Liu, J., Zhang, Q., Zhang, C., Xiang, Y., Zen, K., and Li, L. (2021). Loss of microglial SIRPα promotes synaptic pruning in preclinical models of neurodegeneration. Nat. Commun. 12, 2030. 10.1038/s41467-021-22301-1.

25. De Marchis, S., Fasolo, A., and Puche, A.C. (2004). Subventricular zone-derived neuronal progenitors migrate into the subcortical forebrain of postnatal mice. J. Comp. Neurol. 476, 290–300. 10.1002/cne.20217.

26. Bordiuk, O.L., Smith, K., Morin, P.J., and Semënov, M.V. (2014). Cell proliferation and neurogenesis in adult mouse brain. PloS One 9, e111453. 10.1371/journal.pone.0111453.

27. Ribeiro Xavier, A.L., Kress, B.T., Goldman, S.A., Lacerda de Menezes, J.R., and Nedergaard, M. (2015). A Distinct Population of Microglia Supports Adult Neurogenesis in the Subventricular Zone. J. Neurosci. Off. J. Soc. Neurosci. 35, 11848–11861. 10.1523/JNEUROSCI.1217-15.2015.

28. Renz, P., Surbek, D., Haesler, V., Tscherrig, V., Huang, E.J., Chavali, M., Liddelow, S., Rowitch, D., Schoeberlein, A., and Lutz, A.B. (2022). Silencing neuroinflammatory reactive astrocyte activating factors ameliorates disease outcomes in perinatal white matter injury. Preprint at bioRxiv, 10.1101/2022.12.19.521083 10.1101/2022.12.19.521083.

29. Back, S.A., and Rosenberg, P.A. (2014). Pathophysiology of glia in perinatal white matter injury. Glia 62, 1790–1815. 10.1002/glia.22658.

30. Pfisterer, U., and Khodosevich, K. (2017). Neuronal survival in the brain: neuron type-specific mechanisms. Cell Death Dis. 8, e2643. 10.1038/cddis.2017.64.

31. Castillo-Ruiz, A., Hite, T.A., Yakout, D.W., Rosen, T.J., and Forger, N.G. (2020). Does Birth Trigger Cell Death in the Developing Brain? eNeuro 7, ENEURO.0517-19.2020. 10.1523/ENEURO.0517-19.2020.

32. Dries, R., Zhu, Q., Dong, R., Eng, C.-H.L., Li, H., Liu, K., Fu, Y., Zhao, T., Sarkar, A., Bao, F., et al. (2021). Giotto: a toolbox for integrative analysis and visualization of spatial expression data. Genome Biol. 22, 78. 10.1186/s13059-021-02286-2.

33. Palla, G., Spitzer, H., Klein, M., Fischer, D., Schaar, A.C., Kuemmerle, L.B., Rybakov, S., Ibarra, I.L., Holmberg, O., Virshup, I., et al. (2022). Squidpy: a scalable framework for spatial omics analysis. Nat. Methods 19, 171–178. 10.1038/s41592-021-01358-2.

34. Miyamoto, N., Maki, T., Shindo, A., Liang, A.C., Maeda, M., Egawa, N., Itoh, K., Lo, E.K., Lok, J., Ihara, M., et al. (2015). Astrocytes Promote Oligodendrogenesis after White Matter Damage via Brain-Derived Neurotrophic Factor. J. Neurosci. Off. J. Soc. Neurosci. 35, 14002–14008. 10.1523/JNEUROSCI.1592-15.2015.

35. Koufi, F.-D., Neri, I., Ramazzotti, G., Rusciano, I., Mongiorgi, S., Marvi, M.V., Fazio, A., Shin, M., Kosodo, Y., Cani, I., et al. (2023). Lamin B1 as a key modulator of the developing and aging brain. Front. Cell. Neurosci. 17, 1263310. 10.3389/fncel.2023.1263310.

36. Makwana, M., Jones, L.L., Cuthill, D., Heuer, H., Bohatschek, M., Hristova, M., Friedrichsen, S., Ormsby, I., Bueringer, D., Koppius, A., et al. (2007). Endogenous transforming growth factor beta 1 suppresses inflammation and promotes survival in adult CNS. J. Neurosci. Off. J. Soc. Neurosci. 27, 11201–11213. 10.1523/JNEUROSCI.2255-07.2007.

37. De Lucia, C., Rinchon, A., Olmos-Alonso, A., Riecken, K., Fehse, B., Boche, D., Perry, V.H., and Gomez-Nicola, D. (2016). Microglia regulate hippocampal neurogenesis during chronic neurodegeneration. Brain. Behav. Immun. 55, 179–190. 10.1016/j.bbi.2015.11.001.

38. Battista, D., Ferrari, C.C., Gage, F.H., and Pitossi, F.J. (2006). Neurogenic niche modulation by activated microglia: transforming growth factor beta increases neurogenesis in the adult dentate gyrus. Eur. J. Neurosci. 23, 83–93. 10.1111/j.1460-9568.2005.04539.x.

39. Binder, M.D., Cate, H.S., Prieto, A.L., Kemper, D., Butzkueven, H., Gresle, M.M., Cipriani, T., Jokubaitis, V.G., Carmeliet, P., and Kilpatrick, T.J. (2008). Gas6 deficiency increases oligodendrocyte loss and microglial activation in response to cuprizone-induced demyelination. J. Neurosci. Off. J. Soc. Neurosci. 28, 5195–5206. 10.1523/JNEUROSCI.1180-08.2008.

40. Binder, M.D., Xiao, J., Kemper, D., Ma, G.Z.M., Murray, S.S., and Kilpatrick, T.J. (2011). Gas6 increases myelination by oligodendrocytes and its deficiency delays recovery following cuprizone-induced demyelination. PloS One 6, e17727. 10.1371/journal.pone.0017727.

41. Fujioka, T., Kaneko, N., Ajioka, I., Nakaguchi, K., Omata, T., Ohba, H., Fässler, R., García-Verdugo, J.M., Sekiguchi, K., Matsukawa, N., et al. (2017). β1 integrin signaling promotes neuronal migration along vascular scaffolds in the post-stroke brain. EBioMedicine 16, 195–203. 10.1016/j.ebiom.2017.01.005.

42. Bellver-Landete, V., Bretheau, F., Mailhot, B., Vallières, N., Lessard, M., Janelle, M.-E., Vernoux, N., Tremblay, M.-È., Fuehrmann, T., Shoichet, M.S., et al. (2019). Microglia are an essential component of the neuroprotective scar that forms after spinal cord injury. Nat. Commun. 10, 518. 10.1038/s41467-019-08446-0.

43. Zhou, Z.-L., Xie, H., Tian, X.-B., Xu, H.-L., Li, W., Yao, S., and Zhang, H. (2023). Microglial depletion impairs glial scar formation and aggravates inflammation partly by inhibiting STAT3 phosphorylation in astrocytes after spinal cord injury. Neural Regen. Res. 18, 1325–1331. 10.4103/1673-5374.357912.

44. Gerber, Y.N., Saint-Martin, G.P., Bringuier, C.M., Bartolami, S., Goze-Bac, C., Noristani, H.N., and Perrin, F.E. (2018). CSF1R Inhibition Reduces Microglia Proliferation, Promotes Tissue Preservation and Improves Motor Recovery After Spinal Cord Injury. Front. Cell. Neurosci. 12, 368. 10.3389/fncel.2018.00368.

45. Hammond, T.R., McEllin, B., Morton, P.D., Raymond, M., Dupree, J., and Gallo, V. (2015). Endothelin-B Receptor Activation in Astrocytes Regulates the Rate of Oligodendrocyte Regeneration during Remyelination. Cell Rep. 13, 2090–2097. 10.1016/j.celrep.2015.11.002.

46. Akhmetshina, A., Palumbo, K., Dees, C., Bergmann, C., Venalis, P., Zerr, P., Horn, A., Kireva, T., Beyer, C., Zwerina, J., et al. (2012). Activation of canonical Wnt signalling is required for TGF-β-mediated fibrosis. Nat. Commun. 3, 735. 10.1038/ncomms1734.

47. Argaw, A.T., Asp, L., Zhang, J., Navrazhina, K., Pham, T., Mariani, J.N., Mahase, S., Dutta, D.J., Seto, J., Kramer, E.G., et al. (2012). Astrocyte-derived VEGF-A drives blood-brain barrier disruption in CNS inflammatory disease. J. Clin. Invest. 122, 2454–2468. 10.1172/JCI60842.

48. Klimaschewski, L., and Claus, P. (2021). Fibroblast Growth Factor Signalling in the Diseased Nervous System. Mol. Neurobiol. 58, 3884–3902. 10.1007/s12035-021-02367-0.

49. Li, W., Chen, Z., Chin, I., Chen, Z., and Dai, H. (2018). The Role of VE-cadherin in Blood-brain Barrier Integrity Under Central Nervous System Pathological Conditions. Curr. Neuropharmacol. 16, 1375–1384. 10.2174/1570159X16666180222164809.

50. Knox, E.G., Aburto, M.R., Clarke, G., Cryan, J.F., and O’Driscoll, C.M. (2022). The blood-brain barrier in aging and neurodegeneration. Mol. Psychiatry 27, 2659–2673. 10.1038/s41380-022-01511-z.

51. Harboe, M., Torvund-Jensen, J., Kjaer-Sorensen, K., and Laursen, L.S. (2018). Ephrin-A1-EphA4 signaling negatively regulates myelination in the central nervous system. Glia 66, 934–950. 10.1002/glia.23293.

52. Siebert, J.R., Conta Steencken, A., and Osterhout, D.J. (2014). Chondroitin sulfate proteoglycans in the nervous system: inhibitors to repair. BioMed Res. Int. 2014, 845323. 10.1155/2014/845323.

53. Dyck, S.M., and Karimi-Abdolrezaee, S. (2015). Chondroitin sulfate proteoglycans: Key modulators in the developing and pathologic central nervous system. Exp. Neurol. 269, 169–187. 10.1016/j.expneurol.2015.04.006.

54. Luo, F., Wang, J., Zhang, Z., You, Z., Bedolla, A., Okwubido-Williams, F., Huang, L.F., Silver, J., and Luo, Y. (2022). Inhibition of CSPG receptor PTPσ promotes migration of newly born neuroblasts, axonal sprouting, and recovery from stroke. Cell Rep. 40, 111137. 10.1016/j.celrep.2022.111137.

55. Boche, D., and Gordon, M.N. (2022). Diversity of transcriptomic microglial phenotypes in aging and Alzheimer’s disease. Alzheimers Dement. J. Alzheimers Assoc. 18, 360–376. 10.1002/alz.12389.

56. Mercurio, D., Fumagalli, S., Schafer, M.K.-H., Pedragosa, J., Ngassam, L.D.C., Wilhelmi, V., Winterberg, S., Planas, A.M., Weihe, E., and De Simoni, M.-G. (2022). Protein Expression of the Microglial Marker Tmem119 Decreases in Association With Morphological Changes and Location in a Mouse Model of Traumatic Brain Injury. Front. Cell. Neurosci. 16, 820127. 10.3389/fncel.2022.820127.

57. Paolicelli, R.C., Sierra, A., Stevens, B., Tremblay, M.-E., Aguzzi, A., Ajami, B., Amit, I., Audinat, E., Bechmann, I., Bennett, M., et al. (2022). Microglia states and nomenclature: A field at its crossroads. Neuron 110, 3458–3483. 10.1016/j.neuron.2022.10.020.

58. Wahane, S., Zhou, X., Zhou, X., Guo, L., Friedl, M.-S., Kluge, M., Ramakrishnan, A., Shen, L., Friedel, C.C., Zhang, B., et al. (2021). Diversified transcriptional responses of myeloid and glial cells in spinal cord injury shaped by HDAC3 activity. Sci. Adv. 7, eabd8811. 10.1126/sciadv.abd8811.

59. Matusova, Z., Hol, E.M., Pekny, M., Kubista, M., and Valihrach, L. (2023). Reactive astrogliosis in the era of single-cell transcriptomics. Front. Cell. Neurosci. 17, 1173200. 10.3389/fncel.2023.1173200.

60. Liddelow, S.A., Guttenplan, K.A., Clarke, L.E., Bennett, F.C., Bohlen, C.J., Schirmer, L., Bennett, M.L., Münch, A.E., Chung, W.-S., Peterson, T.C., et al. (2017). Neurotoxic reactive astrocytes are induced by activated microglia. Nature 541, 481–487. 10.1038/nature21029.

61. Zamanian, J.L., Xu, L., Foo, L.C., Nouri, N., Zhou, L., Giffard, R.G., and Barres, B.A. (2012). Genomic analysis of reactive astrogliosis. J. Neurosci. Off. J. Soc. Neurosci. 32, 6391–6410. 10.1523/JNEUROSCI.6221-11.2012.

62. Kucukdereli, H., Allen, N.J., Lee, A.T., Feng, A., Ozlu, M.I., Conatser, L.M., Chakraborty, C., Workman, G., Weaver, M., Sage, E.H., et al. (2011). Control of excitatory CNS synaptogenesis by astrocyte-secreted proteins Hevin and SPARC. Proc. Natl. Acad. Sci. U. S. A. 108, E440–449. 10.1073/pnas.1104977108.

63. Floriddia, E.M., Lourenço, T., Zhang, S., van Bruggen, D., Hilscher, M.M., Kukanja, P., Gonçalves Dos Santos, J.P., Altınkök, M., Yokota, C., Llorens-Bobadilla, E., et al. (2020). Distinct oligodendrocyte populations have spatial preference and different responses to spinal cord injury. Nat. Commun. 11, 5860. 10.1038/s41467-020-19453-x.

64. Marques, S., Zeisel, A., Codeluppi, S., van Bruggen, D., Mendanha Falcão, A., Xiao, L., Li, H., Häring, M., Hochgerner, H., Romanov, R.A., et al. (2016). Oligodendrocyte heterogeneity in the mouse juvenile and adult central nervous system. Science 352, 1326–1329. 10.1126/science.aaf6463.

65. Takeuchi, A., Takahashi, Y., Iida, K., Hosokawa, M., Irie, K., Ito, M., Brown, J.B., Ohno, K., Nakashima, K., and Hagiwara, M. (2020). Identification of Qk as a Glial Precursor Cell Marker that Governs the Fate Specification of Neural Stem Cells to a Glial Cell Lineage. Stem Cell Rep. 15, 883–897. 10.1016/j.stemcr.2020.08.010.

66. Ulanska-Poutanen, J., Mieczkowski, J., Zhao, C., Konarzewska, K., Kaza, B., Pohl, H.B., Bugajski, L., Kaminska, B., Franklin, R.J., and Zawadzka, M. (2018). Injury-induced perivascular niche supports alternative differentiation of adult rodent CNS progenitor cells. eLife 7, e30325. 10.7554/eLife.30325.

67. Ridderstad Wollberg, A., Ericsson-Dahlstrand, A., Juréus, A., Ekerot, P., Simon, S., Nilsson, M., Wiklund, S.-J., Berg, A.-L., Ferm, M., Sunnemark, D., et al. (2014). Pharmacological inhibition of the chemokine receptor CX3CR1 attenuates disease in a chronic-relapsing rat model for multiple sclerosis. Proc. Natl. Acad. Sci. U. S. A. 111, 5409–5414. 10.1073/pnas.1316510111.

68. de Almeida, M.M.A., Watson, A.E.S., Bibi, S., Dittmann, N.L., Goodkey, K., Sharafodinzadeh, P., Galleguillos, D., Nakhaei-Nejad, M., Kosaraju, J., Steinberg, N., et al. (2023). Fractalkine enhances oligodendrocyte regeneration and remyelination in a demyelination mouse model. Stem Cell Rep. 18, 519–533. 10.1016/j.stemcr.2022.12.001.

69. Molina-Gonzalez, I., Holloway, R.K., Jiwaji, Z., Dando, O., Kent, S.A., Emelianova, K., Lloyd, A.F., Forbes, L.H., Mahmood, A., Skripuletz, T., et al. (2023). Astrocyte-oligodendrocyte interaction regulates central nervous system regeneration. Nat. Commun. 14, 3372. 10.1038/s41467-023-39046-8.

70. Hensch, T.K. (2005). Critical period plasticity in local cortical circuits. Nat. Rev. Neurosci. 6, 877– 888. 10.1038/nrn1787.

71. Fawcett, J.W., Oohashi, T., and Pizzorusso, T. (2019). The roles of perineuronal nets and the perinodal extracellular matrix in neuronal function. Nat. Rev. Neurosci. 20, 451–465. 10.1038/s41583-019-0196-3.

72. Willis, A., Pratt, J.A., and Morris, B.J. (2022). Enzymatic Degradation of Cortical Perineuronal Nets Reverses GABAergic Interneuron Maturation. Mol. Neurobiol. 59, 2874–2893. 10.1007/s12035-022-02772-z.

73. Reichelt, A.C., Hare, D.J., Bussey, T.J., and Saksida, L.M. (2019). Perineuronal Nets: Plasticity, Protection, and Therapeutic Potential. Trends Neurosci. 42, 458–470. 10.1016/j.tins.2019.04.003.

74. Lensjø, K.K., Lepperød, M.E., Dick, G., Hafting, T., and Fyhn, M. (2017). Removal of Perineuronal Nets Unlocks Juvenile Plasticity Through Network Mechanisms of Decreased Inhibition and Increased Gamma Activity. J. Neurosci. Off. J. Soc. Neurosci. 37, 1269–1283. 10.1523/JNEUROSCI.2504-16.2016.

75. Batchelor, P.E., Tan, S., Wills, T.E., Porritt, M.J., and Howells, D.W. (2008). Comparison of inflammation in the brain and spinal cord following mechanical injury. J. Neurotrauma 25, 1217– 1225. 10.1089/neu.2007.0308.

76. van der Poel, M., Ulas, T., Mizee, M.R., Hsiao, C.-C., Miedema, S.S.M., Adelia, null, Schuurman, K.G., Helder, B., Tas, S.W., Schultze, J.L., et al. (2019). Transcriptional profiling of human microglia reveals grey-white matter heterogeneity and multiple sclerosis-associated changes. Nat. Commun. 10, 1139. 10.1038/s41467-019-08976-7.

77. Schneider, J., and Miller, S.P. (2019). Preterm brain Injury: White matter injury. Handb. Clin. Neurol. 162, 155–172. 10.1016/B978-0-444-64029-1.00007-2.

78. Sathyanesan, A., Kundu, S., Abbah, J., and Gallo, V. (2018). Neonatal brain injury causes cerebellar learning deficits and Purkinje cell dysfunction. Nat. Commun. 9, 3235. 10.1038/s41467-018-05656-w.

79. Chavali, M., Ulloa-Navas, M.J., Pérez-Borredá, P., Garcia-Verdugo, J.M., McQuillen, P.S., Huang, E.J., and Rowitch, D.H. (2020). Wnt-Dependent Oligodendroglial-Endothelial Interactions Regulate White Matter Vascularization and Attenuate Injury. Neuron 108, 1130–1145.e5. 10.1016/j.neuron.2020.09.033.

80. Varela-Nallar, L., Rojas-Abalos, M., Abbott, A.C., Moya, E.A., Iturriaga, R., and Inestrosa, N.C. (2014). Chronic hypoxia induces the activation of the Wnt/β-catenin signaling pathway and stimulates hippocampal neurogenesis in wild-type and APPswe-PS1ΔE9 transgenic mice in vivo. Front. Cell. Neurosci. 8, 17. 10.3389/fncel.2014.00017.

81. Dizon, M.L.V., Maa, T., and Kessler, J.A. (2011). The bone morphogenetic protein antagonist noggin protects white matter after perinatal hypoxia-ischemia. Neurobiol. Dis. 42, 318–326. 10.1016/j.nbd.2011.01.023.

82. Wu, M., Hernandez, M., Shen, S., Sabo, J.K., Kelkar, D., Wang, J., O’Leary, R., Phillips, G.R., Cate, H.S., and Casaccia, P. (2012). Differential modulation of the oligodendrocyte transcriptome by sonic hedgehog and bone morphogenetic protein 4 via opposing effects on histone acetylation. J. Neurosci. Off. J. Soc. Neurosci. 32, 6651–6664. 10.1523/JNEUROSCI.4876-11.2012.

83. Bergles, D.E., and Richardson, W.D. (2015). Oligodendrocyte Development and Plasticity. Cold Spring Harb. Perspect. Biol. 8, a020453. 10.1101/cshperspect.a020453.

84. Furusho, M., Dupree, J.L., Nave, K.-A., and Bansal, R. (2012). Fibroblast growth factor receptor signaling in oligodendrocytes regulates myelin sheath thickness. J. Neurosci. Off. J. Soc. Neurosci. 32, 6631–6641. 10.1523/JNEUROSCI.6005-11.2012.

85. Furusho, M., Roulois, A.J., Franklin, R.J.M., and Bansal, R. (2015). Fibroblast growth factor signaling in oligodendrocyte-lineage cells facilitates recovery of chronically demyelinated lesions but is redundant in acute lesions. Glia 63, 1714–1728. 10.1002/glia.22838.

86. Aguirre, A., Dupree, J.L., Mangin, J.M., and Gallo, V. (2007). A functional role for EGFR signaling in myelination and remyelination. Nat. Neurosci. 10, 990–1002. 10.1038/nn1938.

87. Li, Y., Su, P., Chen, Y., Nie, J., Yuan, T.-F., Wong, A.H., and Liu, F. (2022). The Eph receptor A4 plays a role in demyelination and depression-related behavior. J. Clin. Invest. 132, e152187. 10.1172/JCI152187.

88. Kirby, L., Jin, J., Cardona, J.G., Smith, M.D., Martin, K.A., Wang, J., Strasburger, H., Herbst, L., Alexis, M., Karnell, J., et al. (2019). Oligodendrocyte precursor cells present antigen and are cytotoxic targets in inflammatory demyelination. Nat. Commun. 10, 3887. 10.1038/s41467-019-11638-3.

89. Nicaise, A.M., Wagstaff, L.J., Willis, C.M., Paisie, C., Chandok, H., Robson, P., Fossati, V., Williams, A., and Crocker, S.J. (2019). Cellular senescence in progenitor cells contributes to diminished remyelination potential in progressive multiple sclerosis. Proc. Natl. Acad. Sci. 116, 9030–9039. 10.1073/pnas.1818348116.

90. Safaiyan, S., Kannaiyan, N., Snaidero, N., Brioschi, S., Biber, K., Yona, S., Edinger, A.L., Jung, S., Rossner, M.J., and Simons, M. (2016). Age-related myelin degradation burdens the clearance function of microglia during aging. Nat. Neurosci. 19, 995–998. 10.1038/nn.4325.

91. Courtois-Cox, S., Jones, S.L., and Cichowski, K. (2008). Many roads lead to oncogene-induced senescence. Oncogene 27, 2801–2809. 10.1038/sj.onc.1210950.

92. Finak, G., McDavid, A., Yajima, M., Deng, J., Gersuk, V., Shalek, A.K., Slichter, C.K., Miller, H.W., McElrath, M.J., Prlic, M., et al. (2015). MAST: a flexible statistical framework for assessing transcriptional changes and characterizing heterogeneity in single-cell RNA sequencing data. Genome Biol. 16, 278. 10.1186/s13059-015-0844-5.

93. Jin, S., Plikus, M.V., and Nie, Q. (2023). CellChat for systematic analysis of cell-cell communication from single-cell and spatially resolved transcriptomics. Preprint at bioRxiv, 10.1101/2023.11.05.565674 10.1101/2023.11.05.565674.

